# Betacoronavirus-specific alternate splicing

**DOI:** 10.1101/2021.07.02.450920

**Authors:** Guy Karlebach, Bruce Aronow, Stephen B. Baylin, Daniel Butler, Jonathan Foox, Shawn Levy, Cem Meydan, Christopher Mozsary, Amanda M Saravia-Butler, Deanne M Taylor, Eve Wurtele, Christopher E Mason, Afshin Beheshti, Peter N Robinson

## Abstract

Viruses can subvert a number of cellular processes in order to block innate antiviral responses, and many viruses interact with cellular splicing machinery. SARS-CoV-2 infection was shown to suppress global mRNA splicing, and at least 10 SARS-CoV-2 proteins bind specifically to one or more human RNAs. Here, we investigate 17 published experimental and clinical datasets related to SARS-CoV-2 infection as well as datasets from the betacoronaviruses SARS-CoV and MERS as well as Streptococcus pneumonia, HCV, Zika virus, Dengue virus, influenza H3N2, and RSV. We show that genes showing differential alternative splicing in SARS-CoV-2 have a similar functional profile to those of SARS-CoV and MERS and affect a diverse set of genes and biological functions, including many closely related to virus biology. Additionally, the differentially spliced transcripts of cells infected by coronaviruses were more likely to undergo intron-retention, contain a pseudouridine modification and a smaller number of exons than differentially spliced transcripts in the control groups. Viral load in clinical COVID-19 samples was correlated with isoform distribution of differentially spliced genes. A significantly higher number of ribosomal genes are affected by DAS and DGE in betacoronavirus samples, and the betacoronavirus differentially spliced genes are depleted for binding sites of RNA-binding proteins. Our results demonstrate characteristic patterns of differential splicing in cells infected by SARS-CoV-2, SARS-CoV, and MERS, potentially modifying a broad range of cellular functions and affecting a diverse set of genes and biological functions.

## Introduction

Coronavirus SARS-CoV-2 emerged in late 2019 as the third human coronavirus identified in the 21st century. Coronavirus disease 2019 (COVID-19) affects diverse organ systems, including the lungs, digestive tract, kidneys, heart, and brain and is associated with extremely heterogeneous manifestations that range from minimal symptoms to significant hypoxia with acute respiratory distress requiring mechanical ventilation [1, 2]. Coronaviruses are large, enveloped, single-stranded RNA viruses found in humans and other mammals and can cause respiratory, gastrointestinal, and neurological disease. In addition to SARS-CoV-2, two other betacoronaviruses associated with severe disease led to global outbreaks in the last two decades: the Severe Acute Respiratory Syndrome (SARS)-associated coronavirus (SARS-CoV) first reported in 2003 and the Middle East Respiratory Syndrome (MERS)-associated coronavirus (MERS-CoV) first reported in 2012 [3].

Viral genomes encode a limited set of genes, and so viruses rely on the host cellular machinery. Viral components can subvert a number of cellular processes in ways that have evolved to block innate antiviral responses. A number of viruses have been shown to interact with cellular splicing machinery. The process of precursor mRNA (pre-mRNA) splicing involves the removal of introns and the precise joining of exons to form mature mRNA molecules. Over 95% of human genes undergo alternative splicing in a developmental, tissue-specific, or signal transduction-dependent manner. Alternative splicing plays important roles in development, disease, and aging [4]. Although some splicing isoforms are produced in the same proportions in all or most cell types, alternative splicing is often regulated by developmental or differential cues or in response to external stimuli [5]. Modulation of splicing has been shown to be an important mechanism in the host’s response to viral infection [6, 7, 8]. On the other hand, viruses can alter splicing patterns. For instance, the Dengue virus NS5 protein binds to core components of the U5 snRNP, and incorporates in active spliceosomes and pre-mRNA processing. Dengue-virus induced changes in the isoform abundance of antiviral factors IKBKE (inhibitor of *κ*B kinase *ϵ*) in a way that could facilitate viral replication [9]. *MX1* encodes an antiviral protein that is induced by interferon-*α*/*β* and inhibits the replication of many RNA viruses. Both Dengue virus and Herpes simplex virus -1 (HSV1) induces alternative splicing in *MX1* that supports instead of restricting viral infection [10, 11, 9]. Poliovirus infection can result in cleavage of a component of the macromolecular SMN complex that mediates assembly of U snRNP complexes by aiding the heptameric oligomerization of Sm proteins onto U snRNAs [12]. Influenza A encodes a protein that modulates mRNA splicing to degrade the mRNA that encodes RIG-I, which encodes a protein that detect the presence of viral RNAs [13].

At least 10 SARS-CoV-2 proteins show specific binding to one or more human RNAs, including 6 structural non-coding RNAs and 142 mRNAs. A highly specific interaction was shown between the SARS-CoV-2 NSP16 protein and human U1 and U2 snRNAs, which hybridize to the 5’ splice site and to the branchpoint, respectively [14]. NSP16 is an RNA cap modifying enzyme with methytransferase activity that modifies viral RNAs [15]. It was shown that NSP16 additionally suppresses host mRNA splicing. SARS-CoV-2 infection and transfection with NSP16 result in an increase in intron retention in multiple IFN-responsive genes, thereby suggesting a role in splicing in suppressing the host interferon response to SARS-CoV-2 infection [14]. Another non-structural protein, NSP1, whose homologs in SARS-CoV and MERS-CoV have roles in viral replication, translational inhibition, transcriptional inhibition, mRNA degradation, and cell cycle arrest, was shown to contribute to global inhibition of host protein translation and manipulating translation [16, 17, 18]; in contrast, the translation of SARS-CoV-2-encoded subgenomic RNAs, which contain a common 5’ leader sequence that is added during negative-strand synthesis is not suppressed. NSP8 and NSP9 bind to the 7SL RNA component of the signal recognition particle (SRP) and interfere with protein trafficking to the cell membrane upon infection. NSP8 and NSP9 appear to be involved in suppression of the interferon response, which is dependent upon the SRP pathway for transport [14].

Here, we present a comprehensive survey of alternative splicing associated with infection by SARS-CoV-2, the two other betacoronaviruses SARS-CoV and MERS, and six other viruses and *Streptococcus pneumonia*, and show associations of betacoronavirus infection with a number of cellular parameters, affecting genes involved in a wide range of biological functions.

## Methods

### RNA-seq data sources

We investigated a range of RNA-seq datasets in which primary cells or cell lines were infected with different pathogens including the three betacoronaviruses as well as the influenza virus H3N2, the bacterium *Streptococcus pneumonia*, Hepatitis C virus, Zika virus, Dengue virus, and Respiratory Syncytial Virus. All these datasets were downloaded from NCBI’s Short Read Archive (SRA) [19]. Infection of cell lines or primary cells with virus is a commonly used experimental system to investigate host dependency factors and host restriction factors [20].

An additional, clinical dataset was analyzed from nasopharyngeal swab specimens (NPSS) from the New York-Presbyterian Hospital-Weill Cornell Medical Center [27]. We will refer to this dataset as SARS-CoV-2-A in the text. Briefly, nasopharyngeal swabs were collected using the BD Universal Viral Transport Media system (Becton, Dickinson and Company, Franklin Lakes, NJ) from symptomatic patients. Total Nucleic Acid (TNA) was extracted using automated nucleic acid extraction on the QIAsymphony and the DSP Virus/Pathogen Mini Kit (Qiagen). RNA isolation and library preparation is fully described in Butler, et al. [27].

### Mapping of RNA-seq data and isoform calling

For each cohort listed in Table 1, RNA-seq data were obtained from the NCBI SRA resource [19]. Except for the nasal swab (NSPP) dataset [27], all datasets were processed using a snakemake pipeline that performs the following steps: samples are downloaded from the SRA, quality-control is performed using fastp [30], alignment to Genome Reference Consortium Human Build 38 version 91 is done using STAR [31], and isoform quantification carried out by RSEM [32].

**Table 1:**
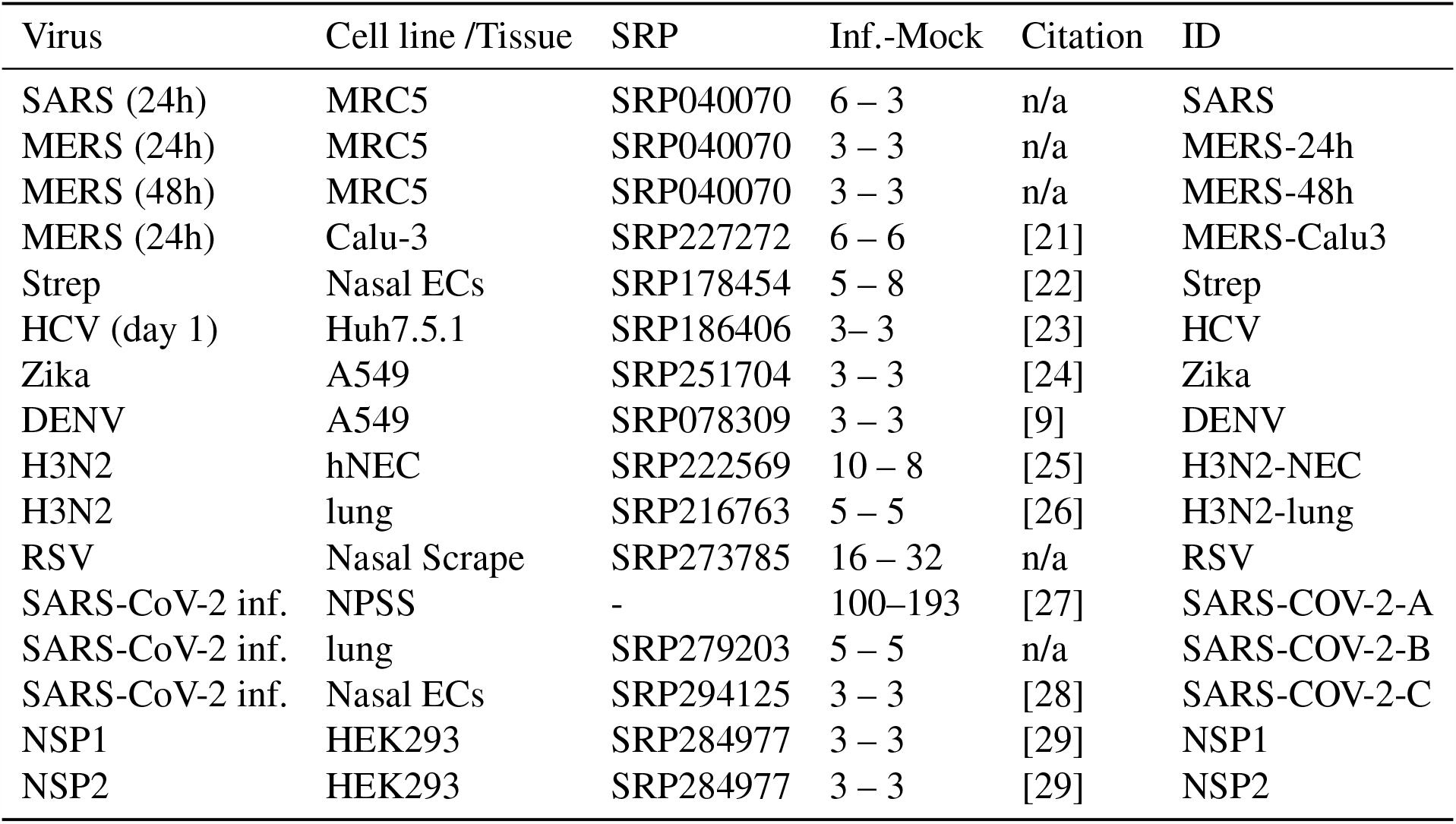
Summary of RNA-seq transcriptional profiling experiments. Data were taken from published experiments in which cell lines were inoculated with an infectious agent and compared to mock inoculations. Datasets were identified in the NCBI short-read archive. Columns: *Inf*.*-Mock*: number of infected/mock samples in an experiment. *ID*: identifier used in this work. Abbreviations: DENV: Dengue virus; ECs: epithelial cells; hNEC: human nasal epithelial cells; MERS: Middle East Respiratory Syndrome Coronavirus; PBMC: Peripheral blood mononuclear cells; SARS: Severe acute respiratory syndrome-associated coronavirus. Strep: *Streptococcus pneumoniae*. RSV: Respiratory Syncytial Virus. The three datasets from SRP040070 were the high multiplicity of infection (MOI) experiments chosen from among a larger set of experiments. NSP1 (NSP2): SARS-CoV-2 NSP1 (NSP2) transfection 24h.

**Table 2:**
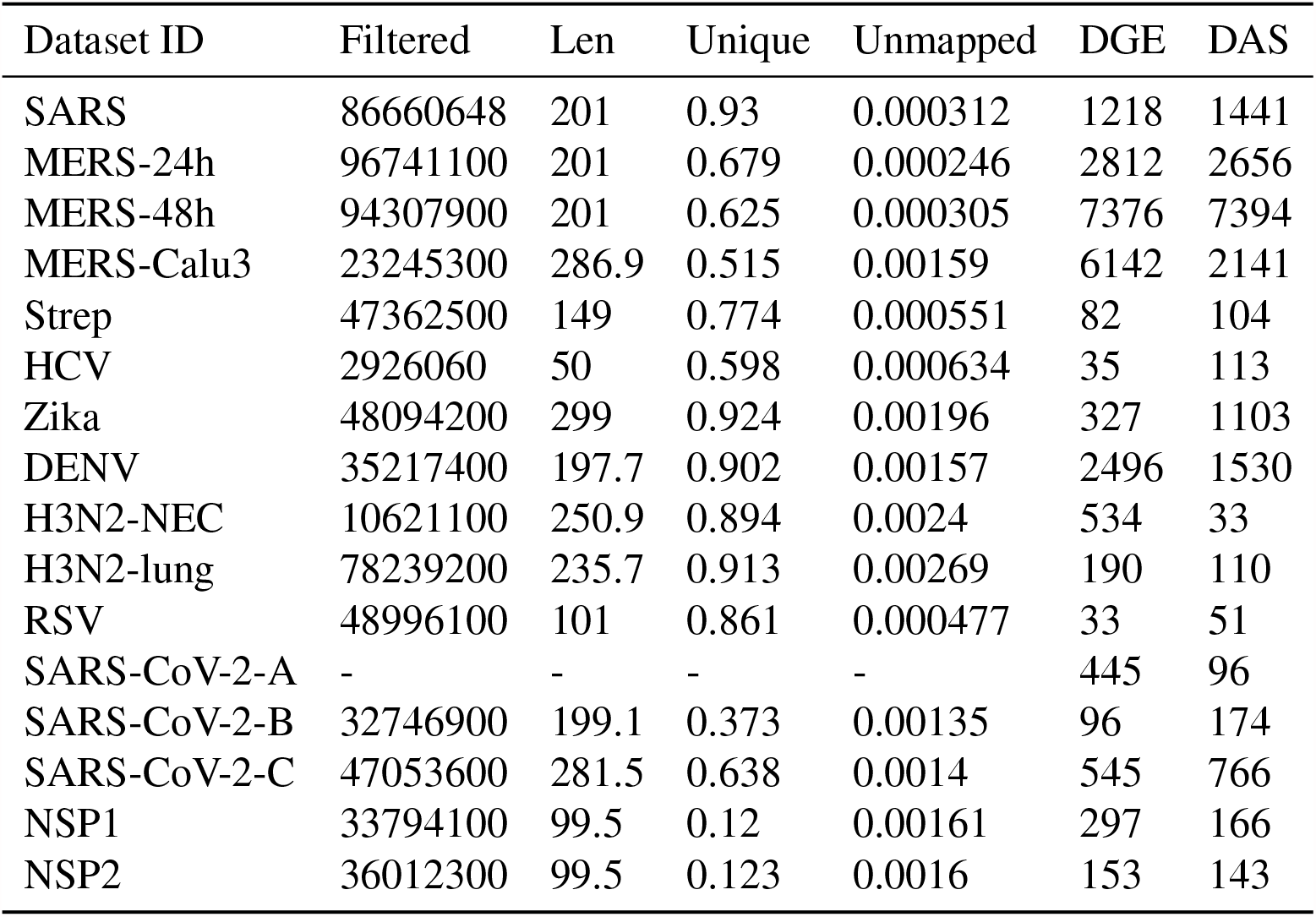
Datasets. Summary of read mapping and HBA-DEALS analysis. Filtered: Mean number of filtered reads; Len: Mean read length; Unique: Mean Fraction Uniquely Mapped Reads; Unmapped: Mean Fraction Unmapped Reads. DGE: Number of differentially expressed genes; DAS: Number of differentially spliced genes. SARS-COV-2-A data were processed using the pipelines developed for COV-iRT [27]. Other datasets were processed as described in the methods.

Raw RNA sequence data from the nasopharyngeal swab samples (SARS-CoV-2-A) were generated as described [27]. Reads classified as human using Kraken2 [33] were processed as described in https://github.com/asaravia-butler/COV-IRT/blob/main/RNAseq/Raw_to_Aligned_Data_Pipeline.md and https://github.com/asaravia-butler/COV-IRT/blob/main/RNAseq/RSEM_Counts_Pipeline.md. First, adapters and low-quality data were trimmed with Trimmomatic (v0.39) [34]. Kraken2-human classified raw and trimmed read quality were evaluated with FastQC (v0.11.9) (https://www.bioinformatics.babraham.ac.uk/projects/fastqc/), and MultiQC (v1.9) [35] was used to generate MultiQC reports. Trimmed reads were split according to sequencing flow cell and lane using gdc-fastq-splitter (v1.0.0) (https://github.com/kmhernan/gdc-fastq-splitter) for subsequent batch effect evaluation. *Homo sapiens* STAR and RSEM references were built using STAR (v2.7.3a) [31] and RSEM (v1.3.1) [32], respectively, with Ensembl release 100 human genome version GRCh38 (`Homo_sapiens.GRCh38.dna.primary_assembly` concatenated with the SARS-CoV-2 Wuhan-Hu-1 SARS-CoV-2 reference genome ASM985889v3, and the following Ensembl gtf annotation file: `Homo_sapiens.GRCh38.100.gtf` concatenated with `Sars_cov_2.ASM985889v3.101.gtf`. rRNA-depleted trimmed reads were aligned to the Homo sapiens and SARS-CoV-2 STAR reference with STAR twopassMode (v2.7.3a) [31]. Transcriptome-aligned reads were quantified using RSEM (v1.3.1) [32] to generate isoform count data.

### Analysis of differential gene expression and splicing

We used HBA-DEALS [36] to calculate the probabilities of differential expression and differential splicing. HBA-DEALS automatically determines the hierarchy of the covariates *α* and *β* - they are either in the same model layer and predict isoform expression, or *β* is in a separate layer that predicts gene expression separately. The choice is made by computing a Bayes factor that compares the two models in a sample of genes, and selecting the model which the majority of Bayes factors favor. For accounting for multiple comparisons we use an Bayesian approach [37]. The prior for *α* and *β* is a mixture of two Dirichlet or two Gaussian components, respectively. The first component has a very large variance and it corresponds to differentially expressed or differentially spliced genes, where little is known about the effect size a-priori. The second component has a very small variance and is centered around 0 for *β* and around 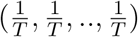 for *α*, where *T* is the number of isoforms. This component corresponds to very small effects that are not biologically meaningful. Since the mixture probabilities are not known in advance, for example we do not know the number of differentially expressed genes, HBA-DEALS infers them from the data. It creates a model that includes all the genes and isoforms in the experiment, and finds the mixture probabilities at the mode of the posterior. The tool stan provides an L-BFGS optimizer which can be accessed via the ‘optimizing’ function in its R interface and which HBA-DEALS uses for this purpose. Once the mixture probabilities for the prior have been obtained, they are set as constant in the prior of each gene, and the full posterior is computed for each gene separately using MCMC sampling, which is executed via the ‘sampling’ function in stan’s R interface. In order to summarize the posteriors of *α* and *β* of each gene into probabilities of differential splicing and differential expression, respectively, HBA-DEALS sums the density of the posterior that falls within a region of parameter values that corresponds to the first component of the prior, i.e. the component that corresponds to differential expression or splicing. The region of parameter values that agrees with the second component of the prior is known as the Region of Practical Equivalence (ROPE) [38]. We place changes of 0.1 in log-expression and 0.2 or less in isoform proportion within the ROPE. After the individual probabilities of effect are calculated, the FDR is estimated as the mean of the probabilities of no effect. In this work we set an FDR threshold of 0.05, and exclude any genes or isoforms with a probability of no-effect greater than 0.25 from the sets of differential genes or isoforms. HBA-DEALS is freely available at https://www.github.com/TheJacksonLaboratory/HBA-DEALS.

### Gene Ontology Analysis

We implemented a version of the Ontologizer [39] analysis code in our Java library phenol (https://github.com/monarch-initiative/phenol). We used the term-for-term GO enrichment analysis with Benjamini-Hochberg correction to select enriched GO terms with FDR≤0.05

### Calculating the Proportion of Retained Intron Isoforms

We used the Ensembl transcript IDs of isoforms included in the study to retrieve the tran-script_biotype field from Biomart [40] for each differentially spliced isoform. The proportion of retained intron isoforms is then the number of ‘retained_intron’ biotypes divided by the total number of biotypes.

### Calculating the Mean Number of Exons of Isoforms

For each transcript, the number of exons in the GTF file Homo_sapiens.GRCh38.91.gtf were counted, and the mean number of exons of differentially spliced isoforms was calculated for each dataset. The SARS-COV-2 nasal swab (NSPP) dataset was aligned to the genome using Homo_sapiens.GRCh38.100.gtf, and therefore for this dataset we used this GTF file for counting the number of exons of each isoform.

## Results

In this study, we have performed an in-depth analysis using publicly available RNA-seq data representing experimental and clinical samples of human cells or tissues infected by a range of viruses and bacteria in order to characterize the biological processes that are affected by changes in gene expression and alternative splicing during SARS-CoV-2, SARS-CoV, and MERS infection and compared them to changes during infection by other viruses and bacteria.

### Disease-associated betacoronaviruses display characteristic functional profile of alternatively spliced genes

In this study, we analyzed RNA-seq datasets representing experimental or clinical studies of SARS-CoV-2, SARS, MERS, four other viruses, and *Streptococcus pneumonia* (Table 1). We reasoned that although differences in experimental design and sample preparation preclude comparisons between individual datasets, a global analysis of profiles of genes and isoforms in each dataset could be used to generate hypotheses about characteristic functional profiles induced by alternative splicing in samples infected with viruses. We therefore performed Gene Ontology (GO) term enrichment analysis on each dataset and created a heat plot for all GO terms in which the BH-corrected *p*-value was less than 0.05 in at least two datasets; a total of 1,044 such terms were identified (Supplemental Figures **S1**-**S2**). Out of 1,044 terms that were enriched in at least two datasets, 1,025 terms showed a high degree of enrichment in at least one betacoronavirus dataset with respect to DAST. 130 of these terms had a median score below threshold (Supplemental Figure **S3**). For display purposes, we chose 14 representative GO terms (Figure 1). Figure 2 displays the -log10 of the adjusted p-value of DAS enrichment in each dataset for two GO terms: *translation* and *RNA binding*.

**Figure 1:**
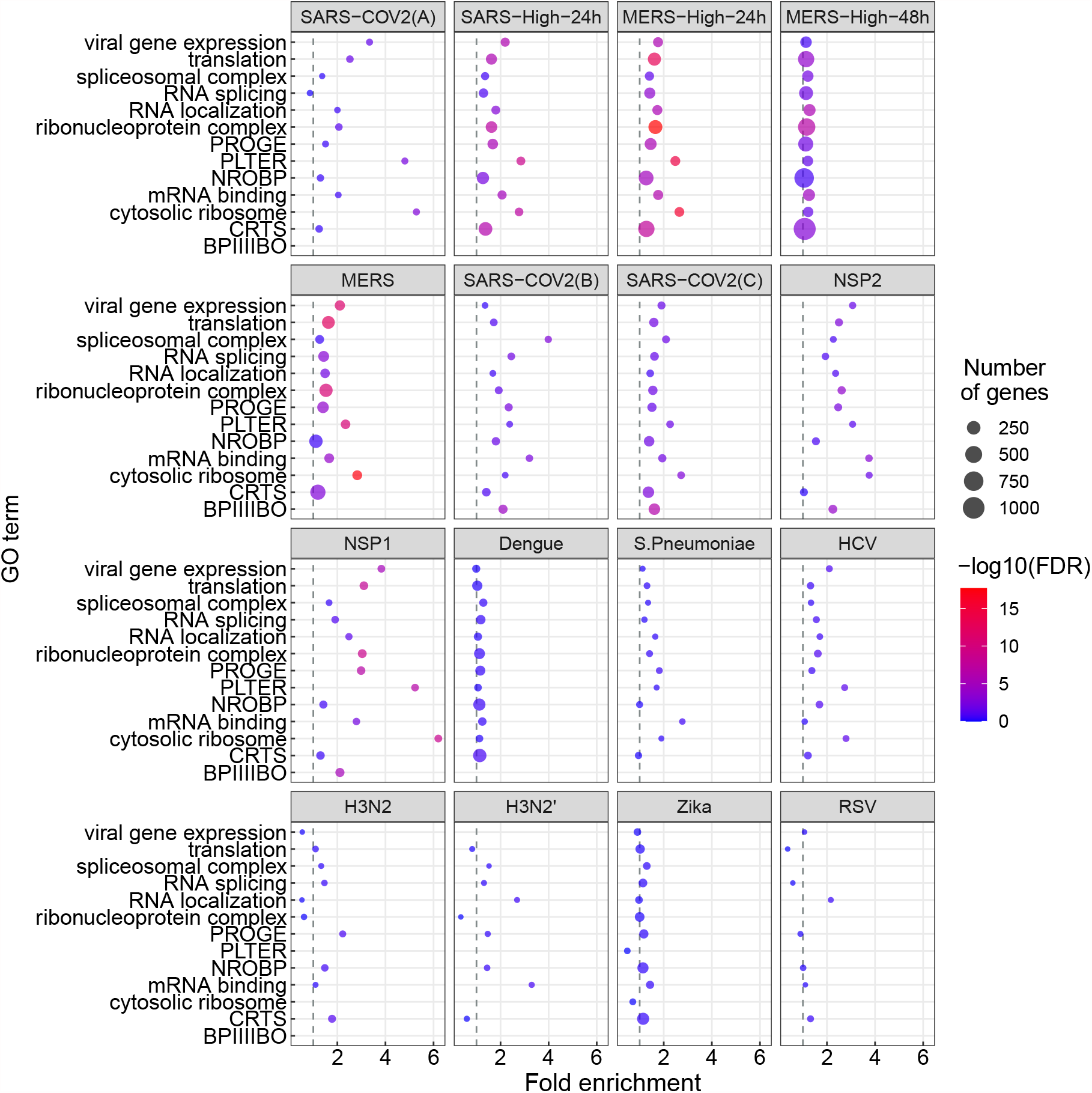
GO enrichment for genes showing differential alternative splicing (DAS). 14 representative GO terms were chosen from a total of 1,044 enriched terms (Supplemental Figure **S1**). The x-axis coordinate corresponds to the fold-change of the size of the GO term in the set of differentially-spliced genes compared to all the genes in the study. The size of the circle corresponds to the number of differentially-spliced genes that belong to the GO term. The color of the circle corresponds to the value of the − log_10_ of the corrected GO term enrichment p-values. Abbreviations. BPIIIIBO: biological process involved in interspecies interaction between organisms; CRTS: cellular response to stress; NROBP: negative regulation of biosynthetic process; PLTER: protein localization to endoplasmic reticulum; PROGE: posttranscriptional regulation of gene expression.

**Figure 2:**
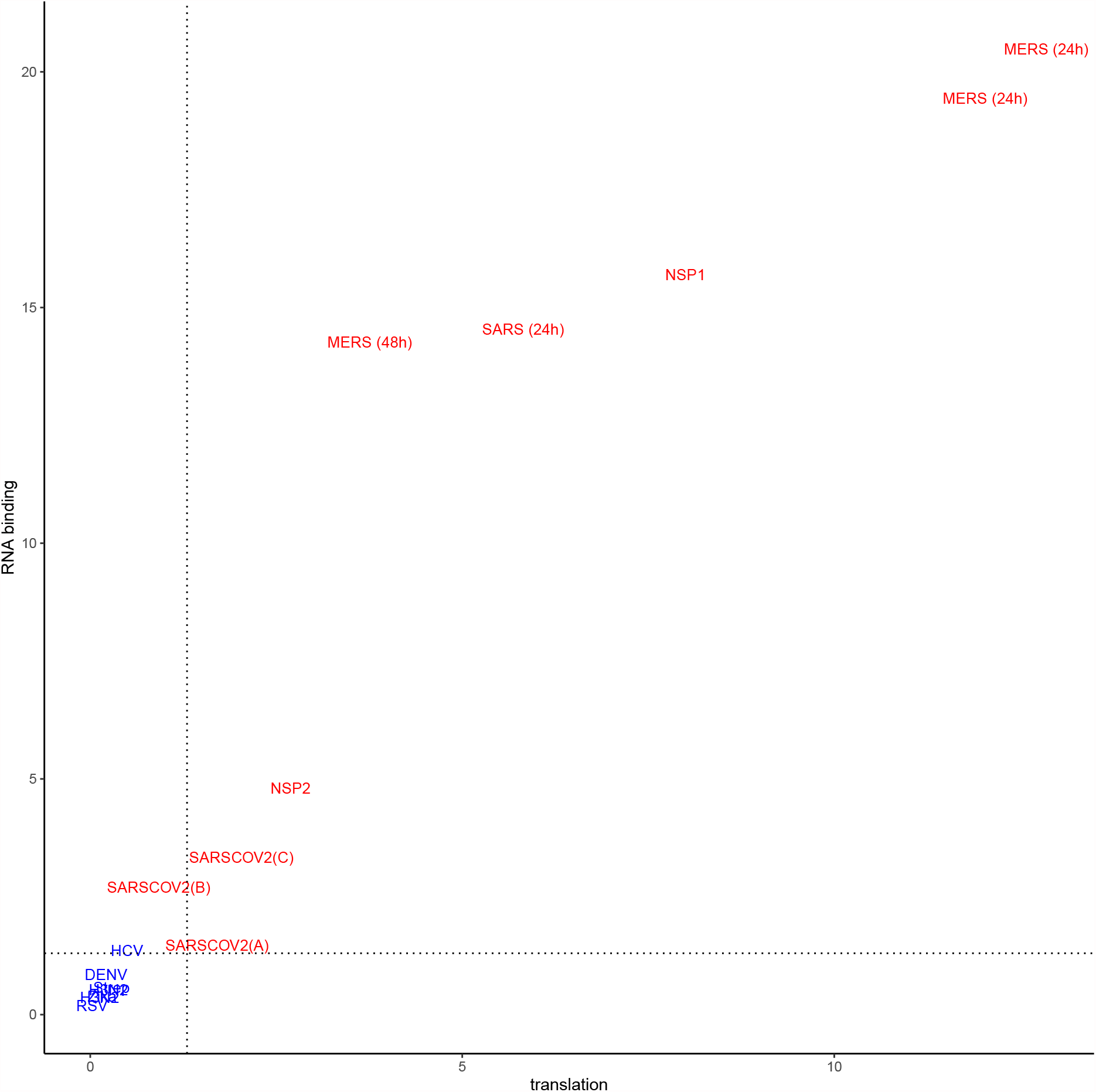
− log_10_(adjusted p value) for DAS enrichment for the GO terms *translation* (GO:0006412) and *RNA binding* (GO:0003723) in each of the datasets analyzed in this work. Betacoronavirus datasets are shown in red, others are shown in blue. The dashed lines correspond to an adjusted p-value of 0.05. Betacoronaviruses have higher enrichment scores for translation and RNA binding, suggesting that differential splicing has a larger impact on these processes. For abbreviations see Table 1.

SARS-CoV-2 infection is reported to disrupt both translation and RNA splicing [14]. We therefore asked if the functional profile of DAS genes is enriched for Gene Ontology (GO) terms related to translation and RNA binding. We found that SARS-CoV-2, NSP1, NSP2, SARS, and MERS datasets showed enrichment for both *translation* (GO:0006412) and *RNA binding* (GO:0003723) and that the degree of enrichment was correlated (Figure 2).

We performed a similar analysis for differentially expressed genes (Figure 3). The terms showing enrichment in multiple betacoronavirus samples included terms directly related to viral infection including *viral gene expression*, as well as biological processes known to be involved in the cellular response to selected viral infections, including *mRNA splicing* and *spliceosomal complex* [41, 42, 43], *protein localization to endoplasmic reticulum* with a possible relation to endoplasmic reticulum stress [44], *cytosolic ribosome* [45], and *translation* [46]. Several of these GO terms were also enriched for differentially expressed genes (Figure 3).

**Figure 3:**
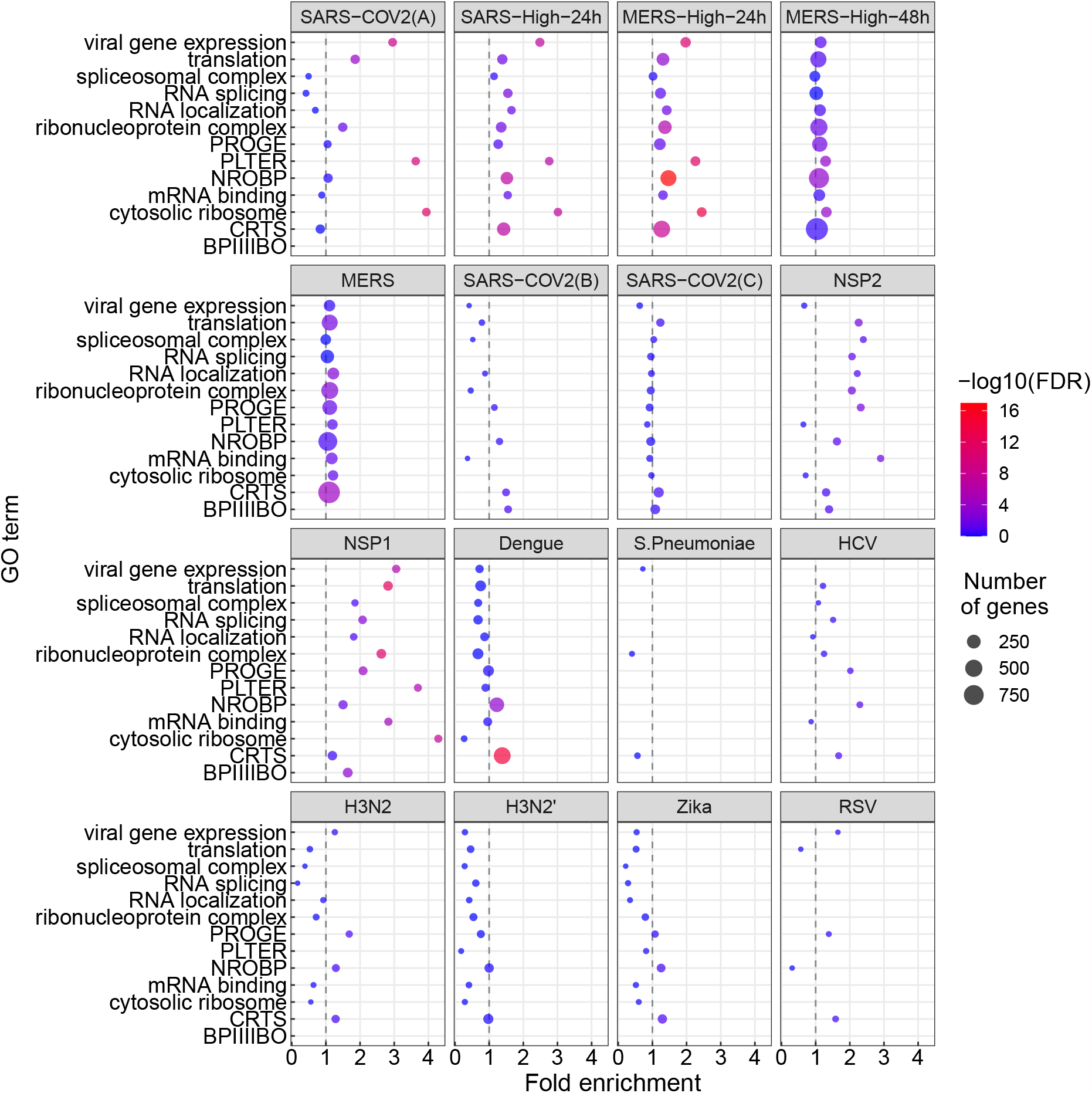
GO enrichment for genes showing differential gene expression (DGE). The GO terms and abbreviations are the same as in Figure 1. The x-axis coordinate corresponds to the fold-change of the size of the GO term in the set of differentially-expressed genes compared to all the genes in the study. The size of the circle corresponds to the number of differentially-expressed genes that belong to the GO term. The color of the circle corresponds to the value of the − log_10_ of the corrected GO term enrichment p-values.

### betacoronavirus samples display more intron retention

SARS-CoV-2 infection was previously shown to disrupt suppress global mRNA splicing [14]. We therefore hypothesized that infection with any of the betacoronaviruses could induce a greater degree of intron retention. We analyzed the mean proportion of intron-retention isoforms among all genes showing DAS and showed a significantly higher degree of intron retention in the betacoronavirus samples (Fig 4A). Additionally, the mean number of exons in isoforms that were significantly differentially spliced in the betacoronaviruses was lower (Fig 4B).

**Figure 4:**
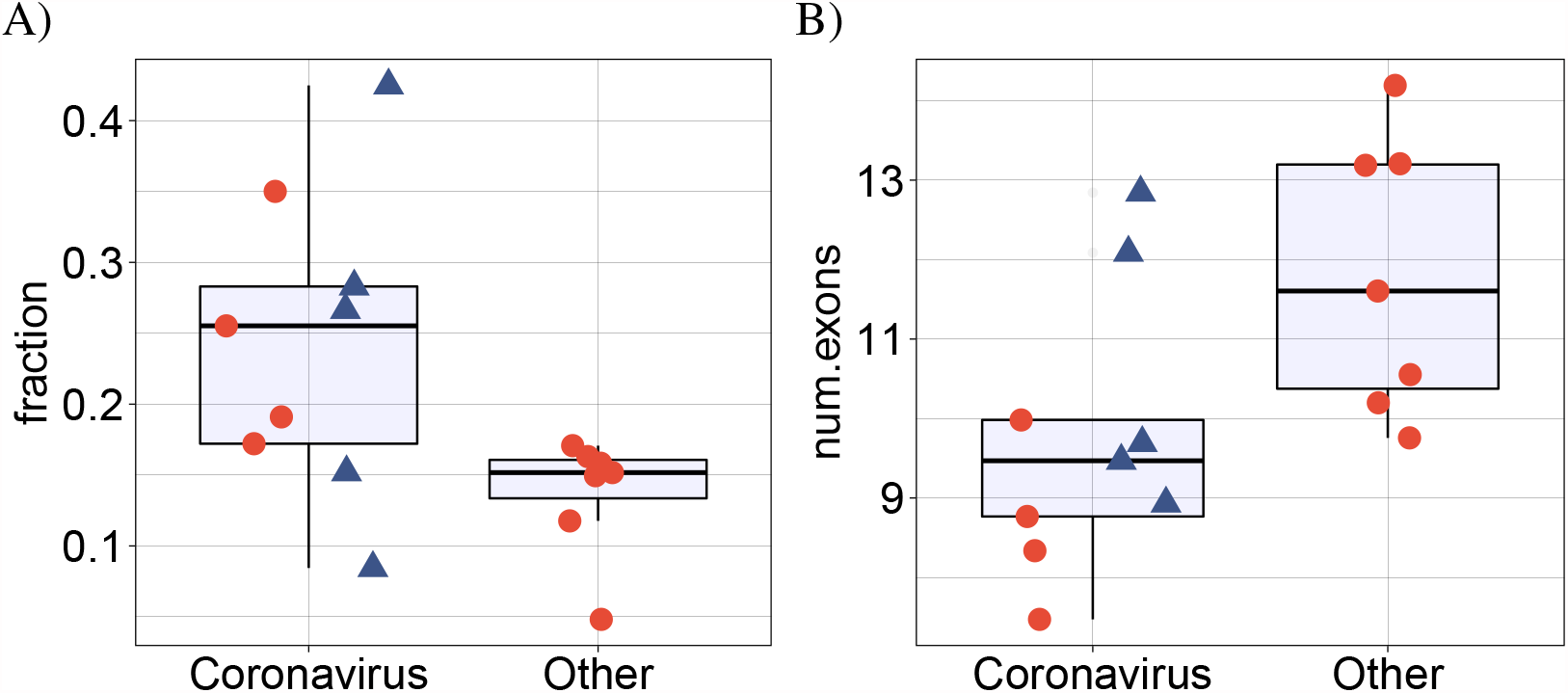
(A) Comparison of the mean proportion of intron retention isoforms in 9 coronavirus samples against the remaining 7 samples for the other pathogens in cyan. *p* = 0.016, Mann-Whitney-test. For each dataset, the mean proportion is calculated as the number of DAS isoforms annotated as retained intron isoforms is divided by the total number of DAS isoforms. (B) Comparison of the mean number of exons in 9 coronavirus samples against the remaining 7 samples for the other pathogens in cyan. *p* = 0.016, Mann-Whitney-test. For each dataset, the mean number of exons is calculated over all DAS isoforms. In both panels, the four SARS-CoV-2 datasets (SARS-CoV-2-A, SARS-CoV-2-B, SARS-CoV-2-C,NSP1 and NSP2) are shown as rectangles.

### More ribosomal genes are affected by DAS and DGE in betacoronavirus samples

Various ribosomal proteins (RPs) have been shown to participate in viral protein biosynthesis and regulate the replication and infection of virus in host cells [45]. We investigated whether the proportion of ribosomal genes displaying DAST and DGE differed between the betacoronavirus and other datasets. We found a significantly higher proportion of ribosomal genes that displayed DAST in the betacoronavirus samples (Fig. 5A).

**Figure 5:**
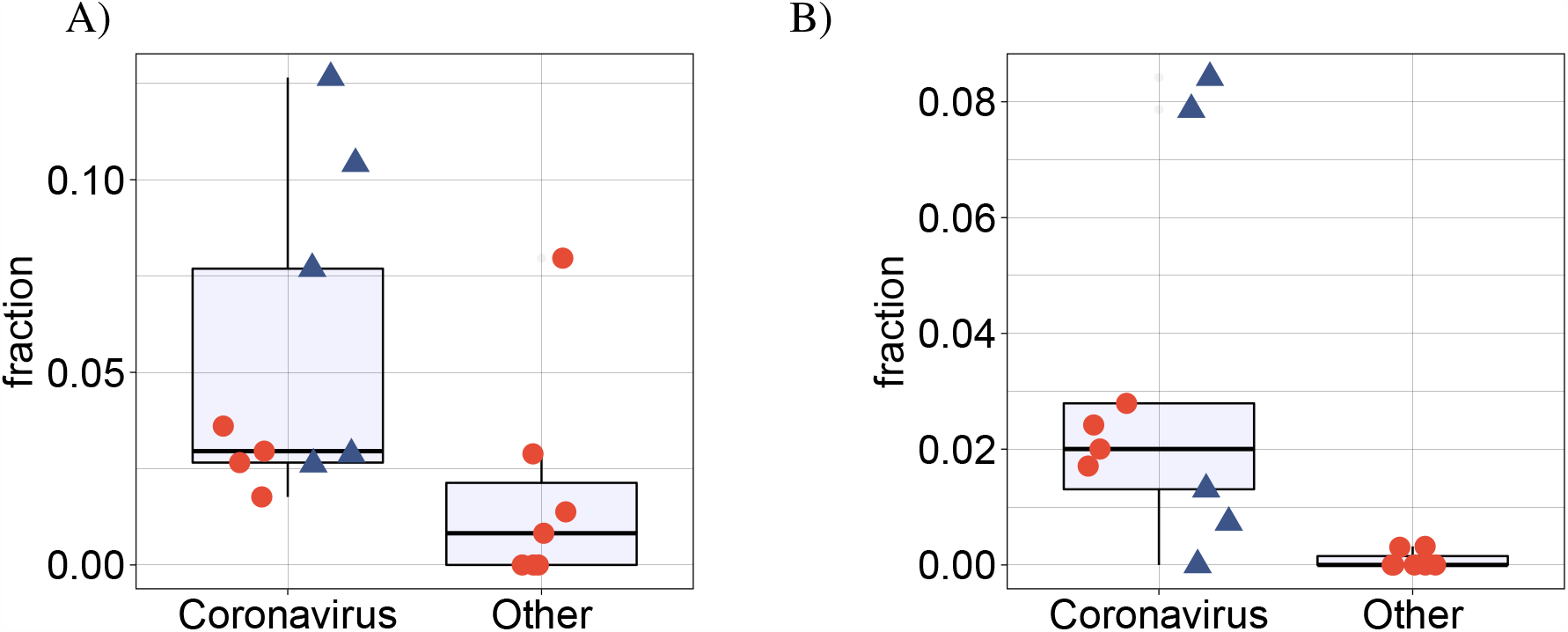
(A) Comparison of the mean proportion of DAS ribosomal genes in 7 coronavirus samples and cells infected with NSP1 or NSP2 against 7 samples infected with other pathogens. *p* = 0.033, Mann-Whitney-test. For each dataset, the mean proportion is calculated as the number of ribosomal genes containing DAS isoforms divided by the total number of genes containing DAS isoforms. (B) Comparison of the mean proportion of differentially expressed ribosomal genes in 7 coronavirus samples and cells infected with NSP1 or NSP2 against 7 samples infected with other pathogens. *p* = 0.004, Mann-Whitney-test. For each dataset, the mean proportion is calculated as the number of ribosomal genes that were differentially expressed divided by the total number of differentially expressed genes. In both panels, the four SARS-CoV-2 datasets (SARS-CoV-2-A, SARS-CoV-2-B, SARS-CoV-2-C, NSP1 and NSP2) are shown as rectangles.

A similar result was obtained for the proportion of ribosomal genes that were differentially expressed (*p* = 0.004, Mann-Whitney-test, Fig. 5B). It has been previously reported that the SARS-CoV-2 NSP1 protein can interfere with translation [47, 14]. Our findings support the conclusion that betacoronavirus infection involves or results in regulatory changes in the transcription of ribosomal genes.

### More betacoronavirus differentially spliced genes are affected by a pseudouridine modification

Pseudouridine is an RNA modification that has been shown to affect splicing [48]. Using the RBM database of RNA modifications [49], we calculated enrichment of RNA modifications in the datasets of this study and compared enrichment in DAS genes of beta-coronaviruses and other pathogens. Figure 6 displays the enrichment score for the different datasets and different modifications, calculated as − *log*_10_(*p*), where *p* is the p-value is obtained using the hypergeometric test. Among the different modifications, Pseudouridine was the modification for which there was the largest difference between the number of betacoronavirus datasets that passed a significance threshold of p-value 0.05 and the number of other pathogen datasets that passed this threshold (Mann-Whitney test p-value 0.003, Fig. 6). This suggests that pseudouridine may be associated with some of the changes in alternative splicing induced during betacoronavirus infection. Other modifications were either enriched in smaller subsets of the betacoronavirus datasets or enriched in both beta-coronaviruses and other datasets, suggesting that the modifications may be related to alternative splicing in general or alternative splicing that is triggered by an immune response.

**Figure 6:**
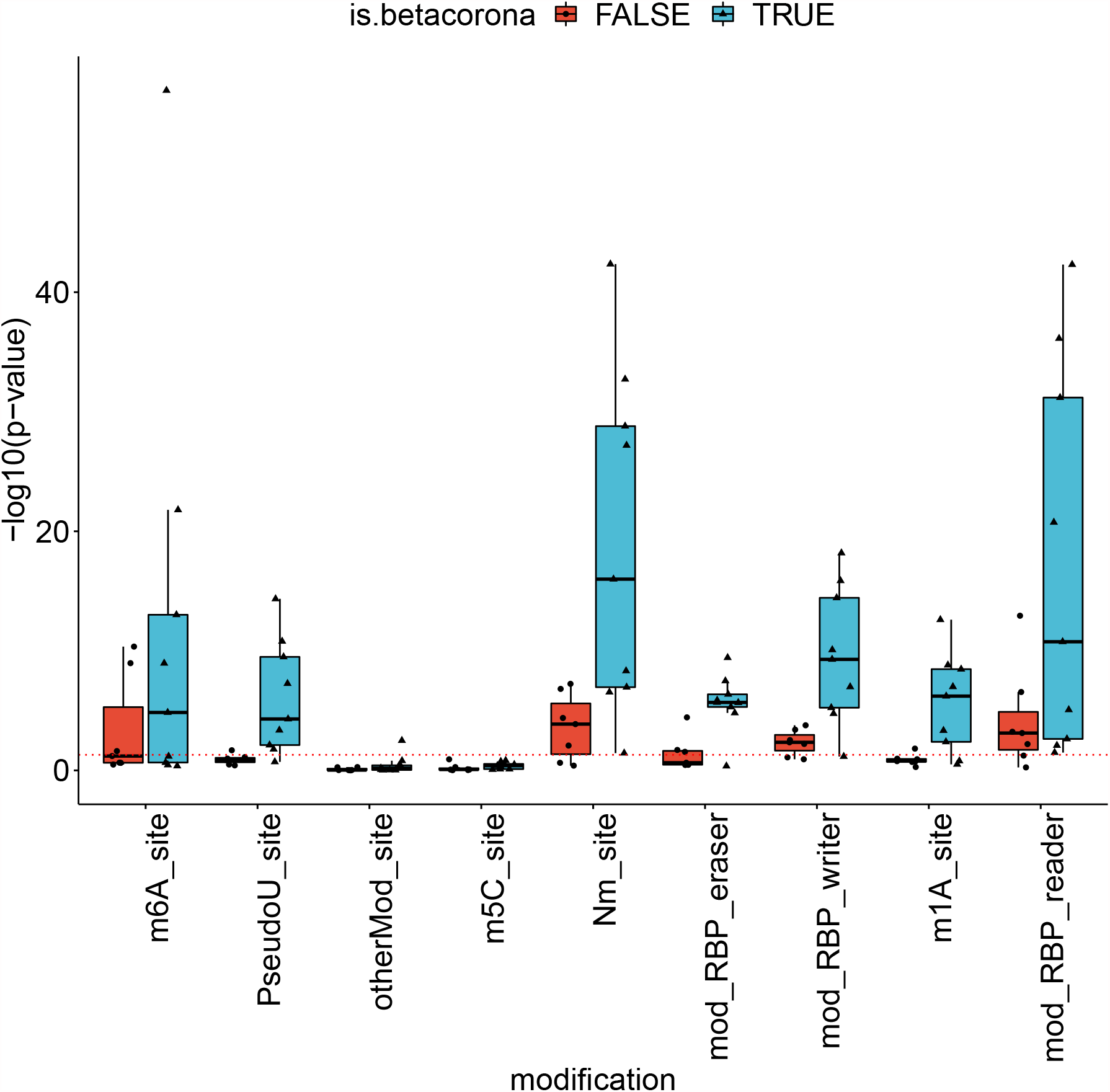
RNA modifications associated with betacoronavirus-enriched alternative splicing. The y-axis corresponds to the enrichment score of the modification in the set of differentially-spliced genes, calculated as − log_10_ of the hypergeometric test p-value. The dashed red line corresponds to a p-value of 0.05. RNA modifications were obtained from the RBM database ([49]). Abbreviations: m6A: N6-methyladenine, PseudoU: pseudouridine, otherMod: other modification, m5c: 5-methylcytosine, Nm: 2^*1*^-O-methylation, RBP eraser: “erasers” of RNA modifications, RBP writer: “writers” of RNA modifications, m1a: N1-methyladenosine, RBP reader: “readers” of RNA modifications.

### Differentially spliced genes of betacoronaviruses are depleted of RBP binding sites

In order to investigate the mechanisms that determine the observed alternative splicing patterns, we calculated the number of binding sites of RNA Binding Proteins (RBPs) in the sets of differentially spliced genes. For this purpose, we downloaded all RBP targets in the human genome from oRNAment [50]. For each RBP binding site, we determined the p-value of observing an identical or higher/lower number of differentially spliced genes among its targets using the hypergeometric test, and corrected the results using Benjamini-Hochberg multiple testing correction. There were 296 RBP binding sites that were enriched for differentially spliced genes over all betacoronavirus datasets, compared to 686 in the other datasets (Figure 7A). Furthermore, there were 687 RBP binding sites whose target sets were depleted of differentially spliced genes in the betacoronavirus datasets, compared to 36 in the other datasets (Figure 7B).

**Figure 7:**
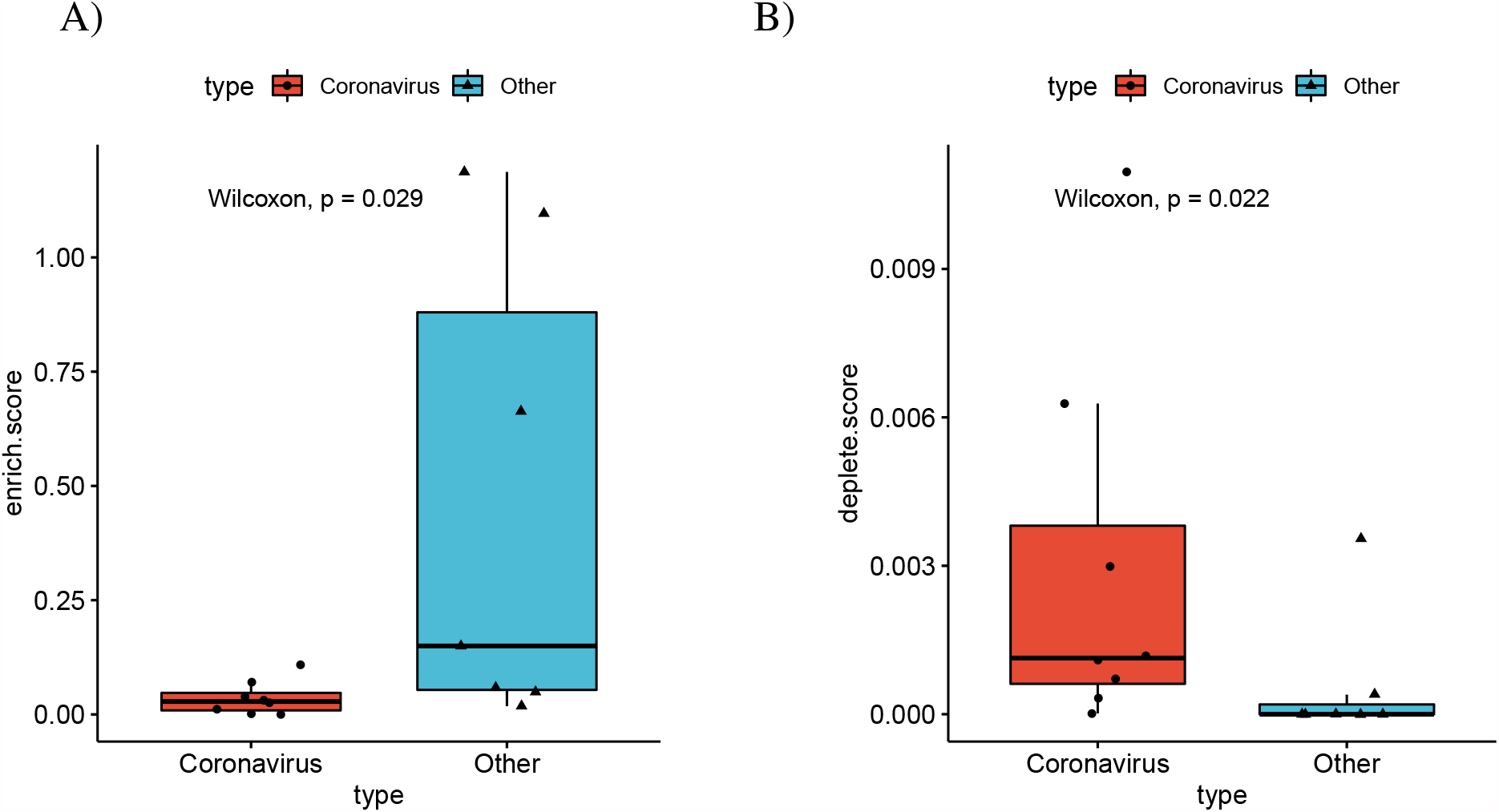
**Differential splicing and enrichment/depletion of RNA Binding Protein Binding Sites**. (A) For each dataset the number of RBP binding sites with enriched targets (adjusted p ≤0.05,hypergeometric test) divided by the number of differentially spliced isoforms are displayed, with separate boxes for betacoronavirus datasets and other datasets(*p* = 0.0289, Mann-Whitney test). RBP binding sites were obtained from the ORNAment database [50](B) Boxplots of the number of RBP binding sites with depleted targets (adjusted p ≤0.05,hypergeometric test) divided by the number of non-differentially spliced isoforms (*p* = 0.0216, Mann-Whitney test).

### Differentially spliced genes of betacoronaviruses are enriched for protein complexes related to ribosome assembly

The CORUM database [51] contains data on experimentally characterized protein complexes. In order to obtain a better understanding of the role of differentially spliced genes in betacoronaviruses, we tested the set of CORUM core complexes for DAS gene enrichment using the hypergeometric test. Setting an FDR threshold of 0.05 using the Benjamini-Hochberg correction, *Nop56p-associated pre-rRNA complex* was enriched in 6 of the 8 betacoronavirus datasets that were downloaded from SRA.

Nop56p is a component of the box C/D small nucleolar ribonucleoprotein complexes that direct 2’-O-methylation of pre-rRNA during its maturation [52]. The Nop56p-associated pre-rRNA complex contains 61 ribosomal proteins including RPL10A, RPL5, RPS3A, RPL4, RPL3, RPL6, RPL22, which were shown to be differentially spliced in our study and are discussed below. Additionally, the CORUM protein complex *Ribosome, cytoplasmic* was enriched in 5 betacoronavirus datasets. Supplemental table **S1** contains all the significant complex enrichments. DAS genes in datasets of other pathogens were not significantly enriched for complexes.

### Viral load is associated with isoform distribution of SARS-CoV-2 DAS genes

We tested the correlation between the fraction of viral RNA and the proportion of counts ofthe different isoform of each differentially spliced gene in the SARS-COV-2-A dataset [27].

For each isoform fractions we performed a Kendall correlation test between the viral load and − log_10_(*p*) for isoforms that belong to DAS genes, and for isoforms of non-DAS genes. Isoforms of DAS genes are more highly correlated with viral load than isoforms of non-DAS genes (Fig. 8). An example for two isoforms of *ADAR* is shown in Supplemental Figure **S4**.

**Figure 8:**
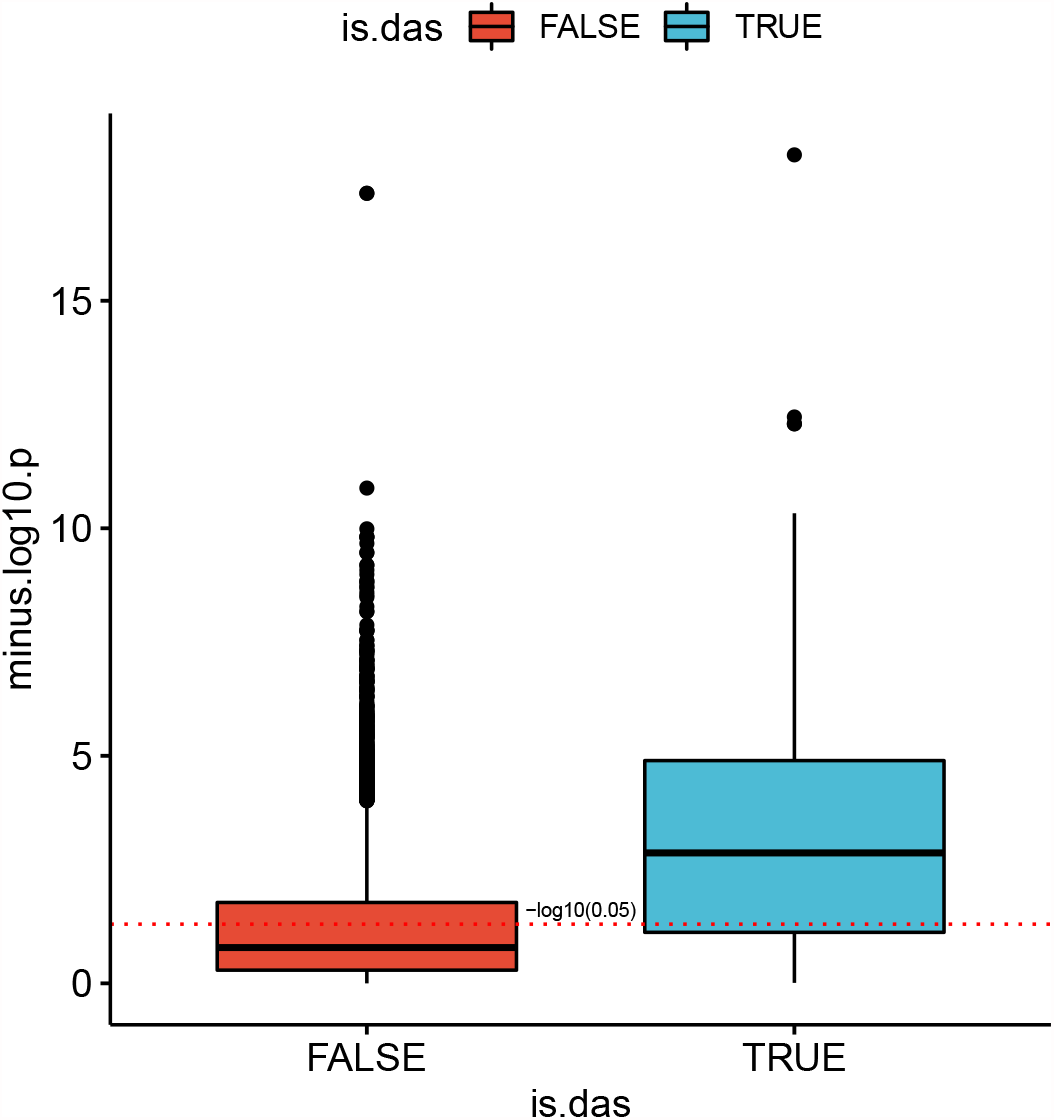
**Isoform distribution and viral load**. The correlation between viral load in each clinical sample and − log_10_(*p*) for isoforms that belong to DAS genes (blue) and for isoforms of non-DAS genes (red) is shown.

For each isoform, the proportion of its counts out of the total number of isoform counts of its corresponding gene was calculated in each sample, and a Kendall correlation test between these values and the fractions of SARS-COV2 RNA was performed. The y-axis corresponds to the − log_10_ of the p-value. The red dashed line corresponds to a p-value of 0.05. The correlation between the raw proportions of differentially-spliced isoforms and suggest that the severity of viral infection as reflected in viral load may be a factor in determining the distribution of patterns of alternative splicing

### Alternative splicing associated with SARS-CoV-2 affects a diverse set of genes and biological functions

It can be challenging to interpret the biological consequences of alternative splicing because experimental characterization of the biological functions of individual isoforms of most genes is not available. However, some of the alternative splicing events detected affected exons or isoforms with known or likely functional roles. Here we present selected alternative splicing events observed in the clinical SARS-CoV-2 nasal swab (NSPP) dataset. Detailed explanations, visualizations, and references are available in Supplemental Figures **S5**–**S10**.

ADARs (adenosine deaminases that act on RNA) target double-stranded RNA (dsRNA) for deamination of adenines into inosines, and act during viral infections to produce hyper-mutation of the viral RNA or to edit host transcripts that modulate the cellular response. ADARs have been show to be involved in coronavirus genome editing [53]. We find that the overall expression of ADAR is increased in COVID-19 patient samples as compared to controls, and in addition, isoforms containing two Z-DNA binding domains are increased whereas an isoform expressing only one such domain is decreased. The shorter isoform with one Z-DNA binding domain is constitutively expressed. The longer isoform is expressed in response to interferon from a different promoter [54]. The smaller isoform is almost exclusively found in the nucleus while the larger is expressed in the cytoplasm [55] (Fig. 9).

**Figure 9:**
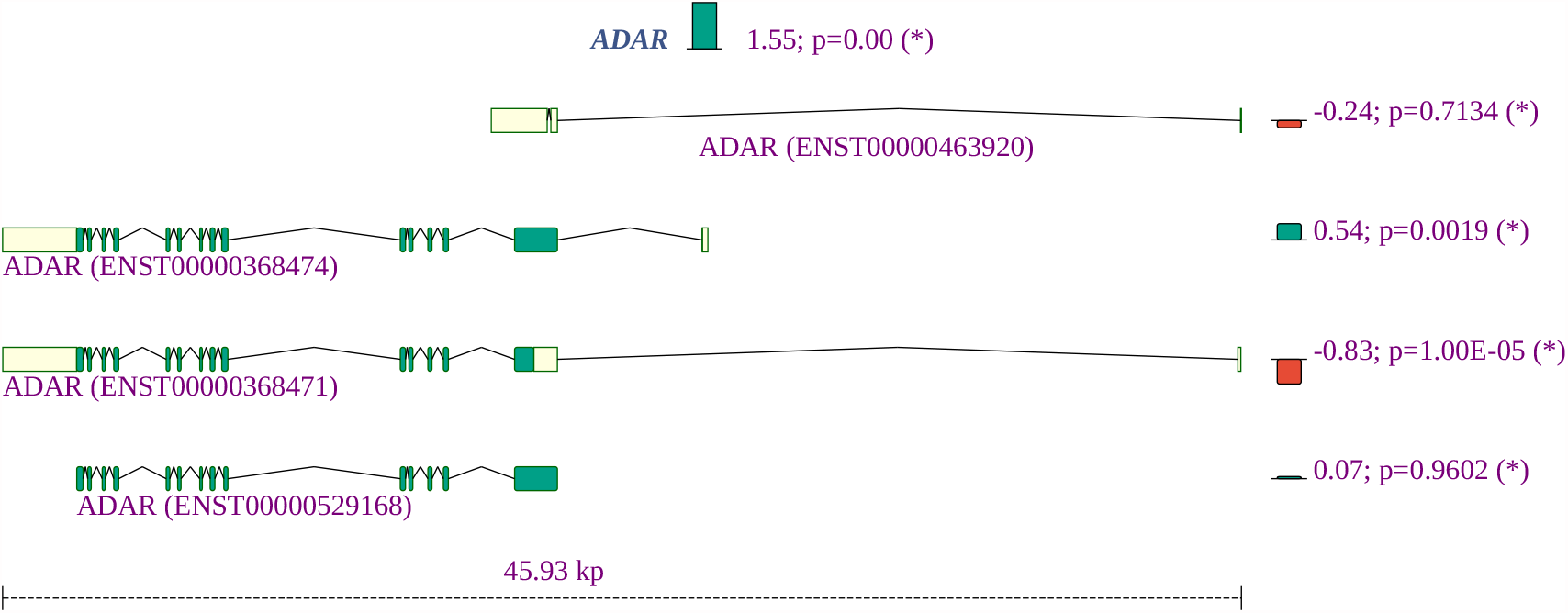
Summary of the gene expression and splicing profile of *ADAR* in COVID-19 patient samples. The expression of *ADAR* was increased by a factor of 2^1.55^ = 3.1. The proportion of isoforms containing two Z-DNA binding domains are increased, whereas an isoform expressing one such domain is decreased.

Adenosine deamination in double-stranded RNAs is mediated by adenosine deaminase acting on RNA (ADAR) enzymes, which can convert adenosine to inosine residues. ADAR enzymes are thought to act on the SARS-CoV-2 genome [53, 56]. Differential splicing of ADAR is likely to affect overall activity of the enzyme and could contribute to the proposed role of ADAR in coronavirus genome editing.

The expression of *NCOA7* was significantly increased. The short isoform of *NCOA7* is inducible by interferon *β* and may play a role in resistance to inflammation-mediated oxidative stress [57]. In our study, the short isoform showed a tendency towards increased expression and the long isoform was significantly underexpressed.

*OAS2*, which was previously shown to be highly overexpressed in lung adenocarcinomal cells infected with SARS-CoV-2 [58], displayed a fold change of 7.11 in the NSPP samples. In addition, a short isoform was a substantially underexpressed compared to the long isoform. The three oligoadenylate synthetases play critical roles in cellular innate antiviral response [59]. The longer isoform of *OAS2* contains two oligoadenylate synthase domains, while the short form contains only one (Supplemental Fig. **S6**).

Strikingly, several ribosomal proteins show both reduced overall expression and a shift from coding to non-coding isoforms, including *RPL10A, RPL22, RPL3,RPL4, RPL5, RPL6, RPS3A*, and *RPS4X* (Supplemental Fig. **S7** shows an example). The small subunit of the ribosome contains one 18S rRNA and about 32 ribosomal proteins (RPs) while, the large 60S subunit consists of 47 ribosomal proteins and one rRNA of 5S, 5.8S, and 28S. This suggests that altered alternative splicing of ribosomal proteins may contribute to the recently described disruption of translation attributed to SARS-CoV-2 infection [14]. In the NSPP dataset, 34 of 409 (8.3%) alternatively spliced genes are annotated to *protein targeting to ER* (GO:0045047), a proportion which is almost 4 times higher than in the population (67/3237; 2.1%). Many ribosomal proteins (RPs) interact with viral mRNA and proteins to participate in viral protein biosynthesis and regulate the replication and infection of virus in host cells [45].

A number of genes were found by HBA-DEALS analysis to be not differentially expressed but to show differential splicing, including *CLSTN1, G3BP1*, and *SMAD3. CLSTN1* encodes calsyntenin, which mediates transport of endosomes along microtubules in neurons as well as mediating intracellular transport of endosomes in HCV-infected cells, thereby contributing to the early stages of the viral replication cycle [60]. A long isoform of *CLSTN1* showed increased expression in the clinical samples (Supplemental Fig. **S8**). *G3BP1* encodes Ras GTPase-activating protein-binding protein 1, an ATP- and magnesium-dependent helicase that plays an essential role in innate immunity that was shown to play an antiviral role against porcine epidemic diarrhea virus, which is a single-stranded, positive-sense RNA virus that belongs to the Coronaviridae [61]. Alternative splicing in clinical samples is a shift to the shorter isoform, which lacks a RNA recognition motif (RRM) domain (Supplemental Fig. **S9**). *SMAD3* encodes an intracellular effector of gene expression responses to TGF-*β*, which can be transcriptionally induced following a number of different viral infections and may promote survival and growth of intracellular pathogens [62]. The SARS-associated coronavirus nucleocapsid protein interacts with Smad3 and modulates transforming growth factor-*β* signaling [63]. In the clinical samples, we noted a shift to a non-coding *SMAD3* isoform (Supplemental Fig. **S10**).

## Discussion

Our study has shown widespread differential alternative splicing associated with SARS-CoV-2, SARS, and MERS infection, affecting genes involved with a characteristic set of functions. We characterized genes showing significant alternative splicing in clinical samples (nasal swabs) of patients with acute COVID-19. Our results provide a catalog of patterns of alternative splicing of potential relevance for understanding the biology of COVID-19 infection.

Although mechanisms differ from virus to virus, the general cycle of infection of a virus involves four major steps: (i) attachment to and entering into a host cell; (ii) replication and transcription of the viral genome followed by translation of viral mRNA; and (iii) assembly into progeny virions; (iv) release of virions from the infected cell. Viruses leverage cellular enzymes to implement these steps. Our analysis identified alternative splicing events potentially affecting each of these functions. For instance, 22 of 175 (12.6%) genes showing alternative splicing in lung tissue infected by SARS-CoV-2 were annotated to *cadherin binding* (GO:0045296), a proportion that is over three times higher than in the population of all genes with at least one read count (239/6435, or 3.7%). One of the differentially spliced genes is *EGFR*. It is known that many viruses usurp EGFR endocytosis or EGFR-mediated signalling for entry into the host cell and other purposes [64]. *IFI16* plays a role in *negative regulation of viral genome replication* (GO:0045071) [65].

However, computational analysis of such datasets remains challenging because limited information is available about specific functions of individual isoforms. Our observation of differential splicing of *IFI16* in SARS-CoV-2 infected lung tissue (SRP279203) is a case in point. *IFI16* is a member of the interferon (IFN)-inducible p200-protein family, all of whose members share a partially-conserved repeat of 200-amino acid residues (also called HIN-200 domain, Prosite:PS50834) in the C-terminus. Additionally, most proteins also share a protein-protein interaction DAPIN domain (prosite:PS50824) in the N-terminus [66, 67]. The IFI16 protein can sense cytosolic as well as nuclear dsDNA and can initiate different innate immune responses. In previous literature, three isoforms of are described, with IFI16A containing all exons, IFI16B not including an exon termed 7a (exon 9 of ENST00000295809.12, exon id ENSE00003664980), and) IFI16C not including exons “7” (exon 8 of ENST00000295809.12, exon id ENSE00003481880) and 7a [68]. IFI16A can thus contain one, two, or three copies of highly conserved 56-amino acids serine-threonine-proline (S/T/P)-rich spacer region. IFI16 was shown to influence both glucocorticoid receptor (GR) transactivation and transrepression via an interaction that was specific to the B isoform [69]. The IFI16 B isoform has been reported to be selectively up-regulated in the inflammatory disease systemic lupus erythematosus [70]. It is not always straightforward to use information like this to interpret findings from RNA-seq studies. Currently, a total of 14 *IFI16* isoforms are registered in Ensembl. We observed differential splicing for isoforms that correspond to isoform A and isoform C in the older literature (Figure 10). It is currently unknown whether isoforms with three (A) or one (C) copy of the spacer region have specific functionality. Our findings suggest that the cellular response to SARS-CoV-2 infection in the lung involves both upregulation of *IFI16* as well as a shift from isoforms with one spacer region to isoforms with three copies. This, and many other similar findings illustrate both the limits of our knowledge of the biological roles of alternative splicing and highlight targets for hypothesis driven research on the functions of differentially spliced isoforms.

**Figure 10:**
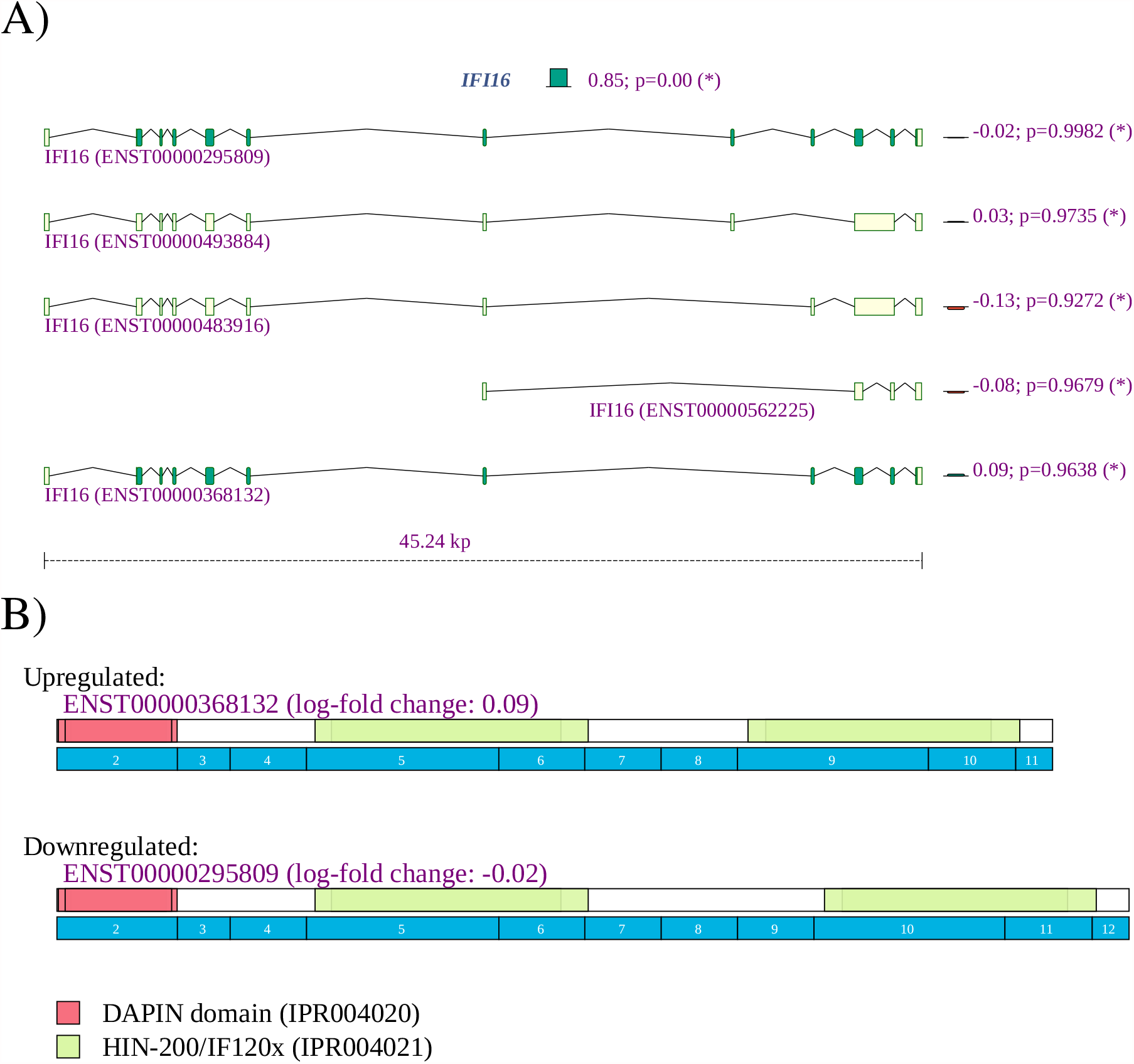
*IFI16* **A)** The expression of *IFI16* is increased with a fold change of 2^2.40^ = 5.23. Reads were mapped to two of the 14 isoforms noted in Ensembl. The expression of isoform ENST00000295809, corresponding to IFI16A, was 2^0.85^ = 1.80 higher in lung samples from COVID-19 patients, and the expression of isoform ENST00000448393, corresponding to IFI16C, was 2^−1.95^ or 3.86 times lower.

Our current data on ADAR must be viewed in the context of the complex, multi-tasking role that this protein plays in immune responses in viral infections and cancer and how the splicing variants we identify factor into these roles. ADAR forms can be important, through interactions with multiple other proteins for controlling levels and directions of immune changes [55, 71]. Key examples include that ADAR has been recently emphasized as important for causing immune suppression and the mechanisms are still being defined [55, 71]. In this regard, ADAR balances self-tolerance and immune activity by modulating canonical antiviral pathways induced by dsRNA [72]. Adenosine to inosine editing or binding of the cytoplasmic ADAR1 isoform p150 and/or the nuclear p110 to dsRNA prevents causes these species to escape cytoplasmic antiviral signaling pathways, via interactions with the RIG-I like receptor-, OAS/RNAseL-, and PKR pathways [55]. This role can be directed towards dsRNA species of viral origin, or, with reduced ADAR levels, endogenous dsRNAs including from inverted Alu repeats and particularly from dsRNA’s residing in mitochondria [55]. In fact, control of inflammasome signaling can be controlled in mitochondria by effects of ADAR on activity of a key protein CMPK2 (PKR). Prior studies have shown that ADAR1 can inhibit viral RNA mediated PKR activation [73] and more recently levels of ADAR have been shown to block the action of CMPK2 in accelerating inflammasome signaling [74]. Further, in viral infections, ADAR is involved in immune antiviral signaling, through regulation of IFN-I production and induction of cellular translation arrest, and apoptosis [73, 74]. These latter activities must be tightly regulated in order to not create an environment that favors virus replication. Finally, it has been proposed that ADAR plays a ‘dual’ protective role against autoinflammatory disease by regulating ‘IFN production’ and the response to IFN. This is especially apparent when ADAR is mutated in a form of childhood “interferonopathy” wherein there is resultant increase in MDA5-mediated ‘IFN production’ in specific cell types such as neuronal lineage, probably explaining resultant severe neuropathology [74].

In addition to providing a comprehensive atlas of genes showing differential splicing related to SARS-CoV-2, we have shown a striking overlap in the functional roles of genes displaying DAS in samples infected with any of the three betacoronaviruses SARS-CoV, SARS-CoV-2, and MERS. We have shown a higher proportion of intron-retention isoforms in cells infected by coronaviruses as compared to a control group of cells infected by *Streptococcus pneumonia*, HCV, Zika virus, Dengue virus, influenza H3N2, and RSV. Speculatively, this could be related to the global suppression of mRNA splicing thought to be due to NSP16 binding to the mRNA recognition domains of the U1 and U2 splicing RNAs [14]. We additionally showed that DAS genes identified in coronavirus-infected samples tend to have a lower number of exons than DAS genes in the control groups. We have no mechanistic explanation for this observation.

In summary, our study provides a comprehensive atlas of genes showing differential alternative splicing associated with infection by SARS-CoV-2, other betacoronaviruses, and a control set of unrelated viruses. Differential alternative splicing occurs in a diverse range of genes that perform a broad range of functions. We found characteristic enrichment of functions related to mRNA binding and splicing, gene expression, and endoplasmic reticulum, among other functions. Our study identified a number of associations that may provide hypotheses for future targeted studies, including increased intron retention, depletion of RBP binding sites in differentially spliced genes, and association of exons affected by alternative splicing with several RNA modifications, and an association of genes affected by alternative splicing with ribosomal complexes.

## Acknowledgments

Support for this work was provided by the Donald A. Roux family fund. A.B. was supported by supplemental funds for COVID-19 research from Translational Research Institute of Space Health through NASA Cooperative Agreement NNX16AO69A (T-0404). We thank New York-Presbyterian Hospital, the Clinical Translational Science Center (ULI TR000457), the Joint Clinical Trials Office, the Core Facilities at Weill Cornell Medicine, the Clinical Laboratories at New York-Presbyterian Hospital, the Scientific Computing Unit (SCU), OneCodex, the XSEDE Supercomputing Resources and the GISAID Initiative curators and submitters. We are grateful for support from Cynthia Polsky, the STARR Foundation (I13-0052) the Vallee Foundation, the WorldQuant Foundation, The Pershing Square Sohn Cancer Research Alliance, Citadel, the National Institutes of Health (R01MH117406, R25EB020393, R01AI151059), the Bill and Melinda Gates Foundation (OPP1151054), the NSF (1840275), the National Center for Advancing Translational Sciences of the National Institutes of Health (UL1TR000457, CTSC), the Intramural Research Program of the National Library of Medicine, NIH, and the Alfred P. Sloan Foundation (G-2015-13964).

## Data and Code Availability Statements

Code for running the pipeline and analysis described in this work is available at https://github.com/TheJacksonLaboratory/under the GNU general public license version 3.

## Declaration of Interests

The authors declare no competing interests.

## Author contributions

G.K and P.N.R. conceived the research. G.K. designed the computational experiments and analyzed the data. D.B., J.F., S.L., C.M, C.M., and C.E.M. collected clinical specimens and performed the RNA-sequencing and bioinformatic processing of the SARS-CoV-2-A dataset. G.K. and P.N.R. worked on data analysis, interpretation, figures, with contributions from all other authors. All authors discussed the results and edited and proofread the manuscript.

## Notes

### Competing Interest Statement

The authors have declared no competing interest.

