## Supplemental material for "Betacoronavirus-specific alternate splicing"

| Complex Name | Number of datasets |
|---|---|
| Nop56p-associated pre-rRNA complex | 6 |
| Ribosome, cytoplasmic | 5 |
| 60S ribosomal subunit, cytoplasmic | 4 |
| 40S ribosomal subunit, cytoplasmic | 3 |
| TRBP containing complex (DICER, RPL7A, EIF6, MOV10 and subunits of the 60S ribosomal particle) | 2 |
| BAF complex | 1 |

**Table S1:** CORUM database core complexes that were enriched for DAS genes in betacoronavirus datasets. The column "Number of datasets" shows the number of betacoronavirus that showed enrichment for the complex.

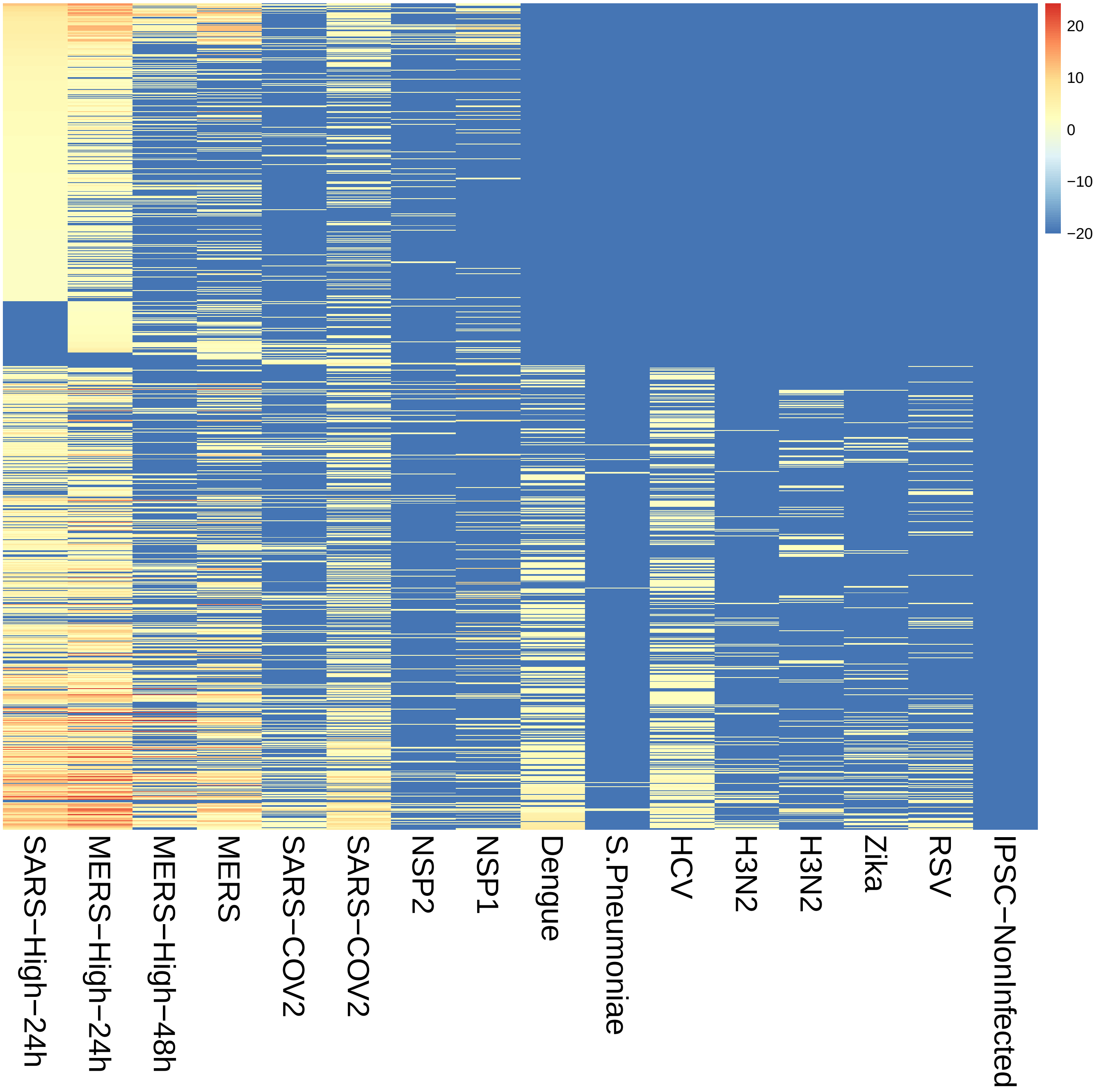


**Figure S1:** Heatplots representing enrichment of Gene Ontology (GO) terms in 16 datasets (See Table 1 of main manuscript). **A)** A total of 1,044 GO terms were significantly enriched in the DAS genes identified in at least two of the 16 investigated datasets. The SARS, SARS-CoV-2, and MERS samples showed the highest number of enriched terms. Each cell of the heat plot represents $-\log_{10} p$, where $p$ is the $p$-value for enrichment of the term in the sample.

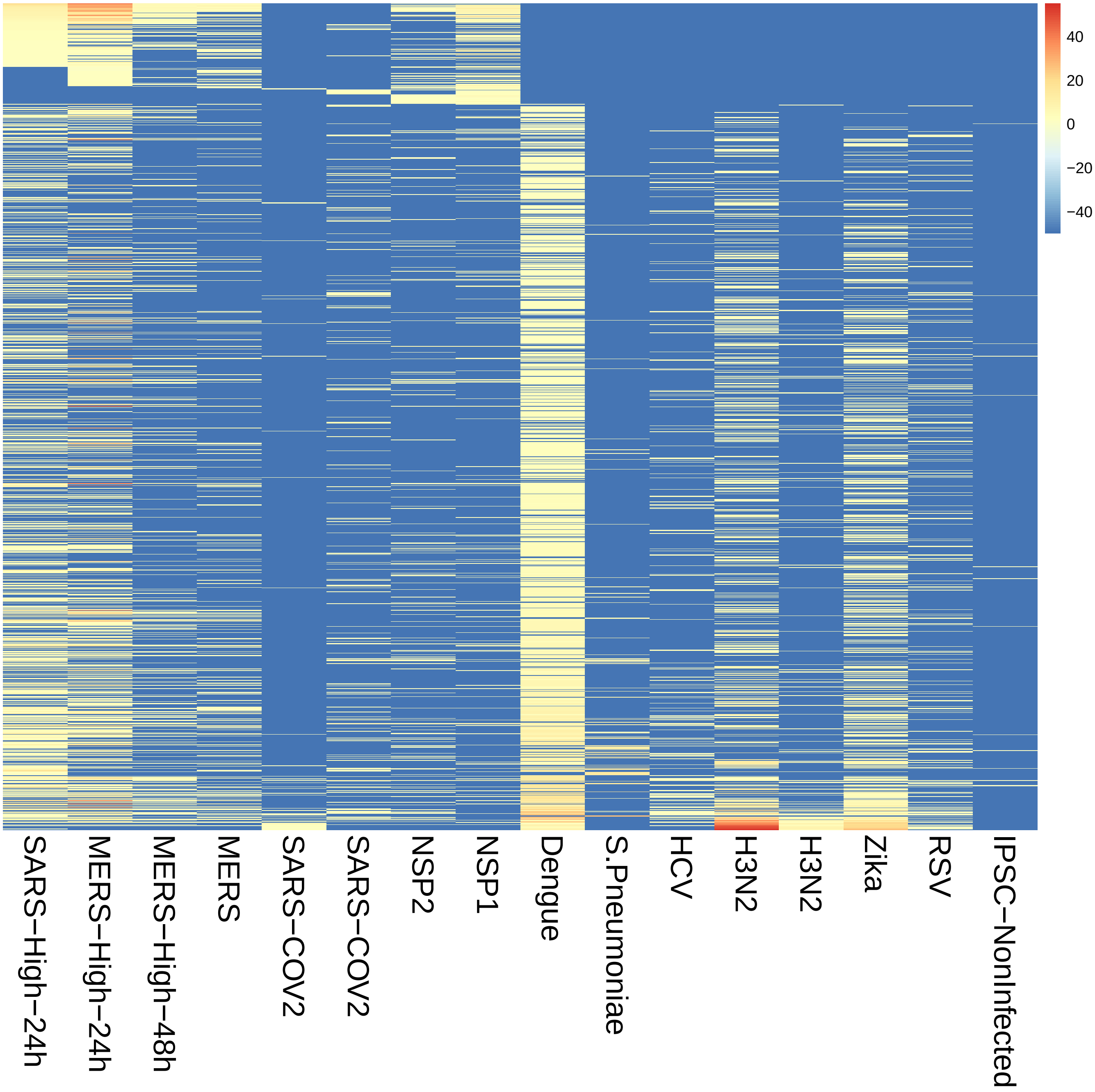


**Figure S2:** Heatplots representing enrichment of Gene Ontology (GO) terms in 16 datasets (See Table 1 of main manuscript). A total of 1,519 GO terms were significantly enriched in the DGE genes identified in at least two of the 16 investigated datasets. Row labels are omitted because of lack of legibility, and details are presented in Figures 1 and 2 of the main manuscript for selected terms. Rows are ordered according to $\max(-\log_{10} p)$ for $p$-values of the non-betacoronavirus datasets.

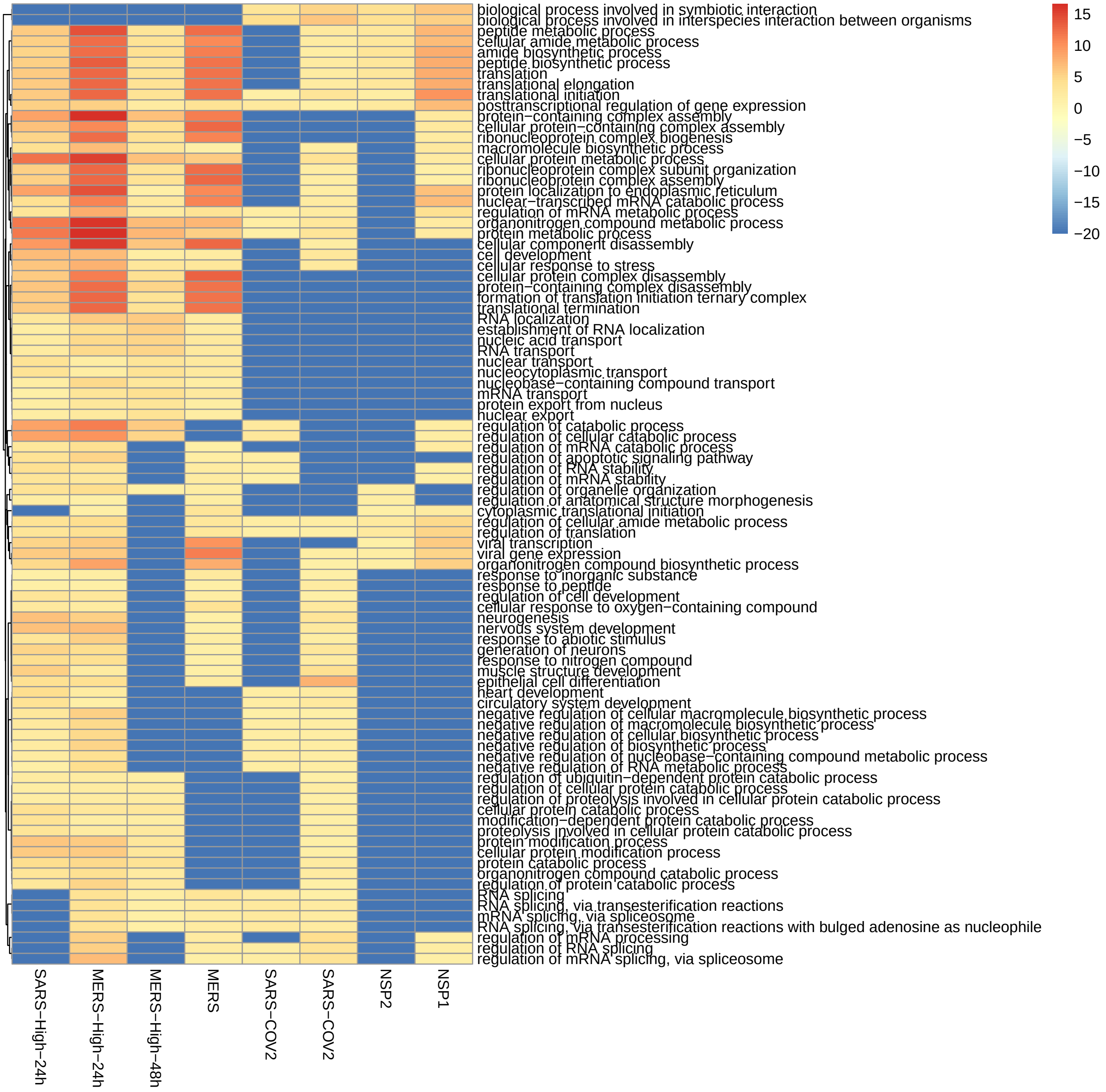


**Figure S3:** 130 of the terms shown in Fig. 1 of the main manuscript had a median score below the threshold, and 90 of them that belong to the Biological Process ontology are shown here. Hierarchical clustering was performed using the R package pheatmap with clustering method ’ward’.

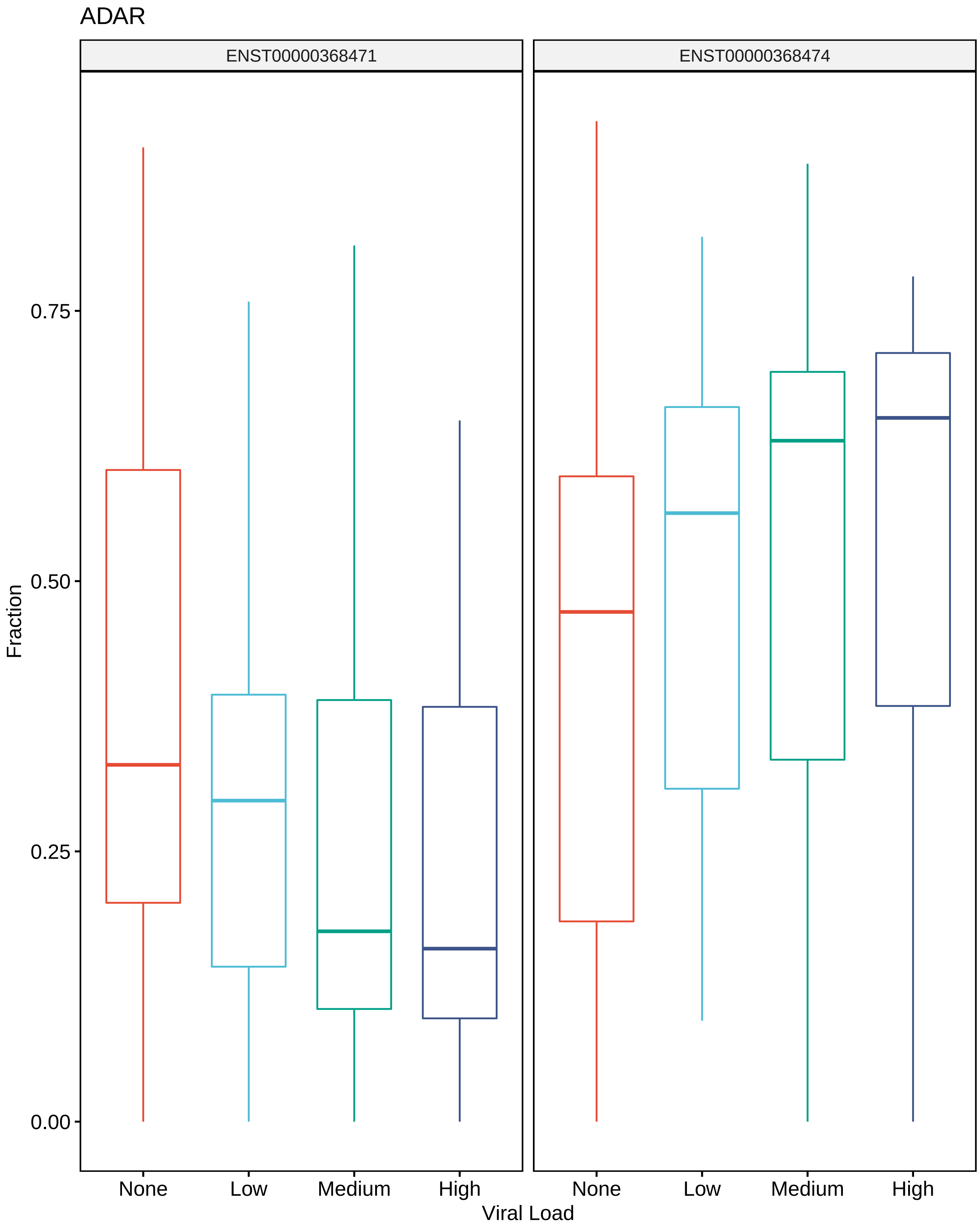


**Figure S4:** Fraction of RSEM counts of ADAR isoforms out of total ADAR RSEM counts, for different viral loads.

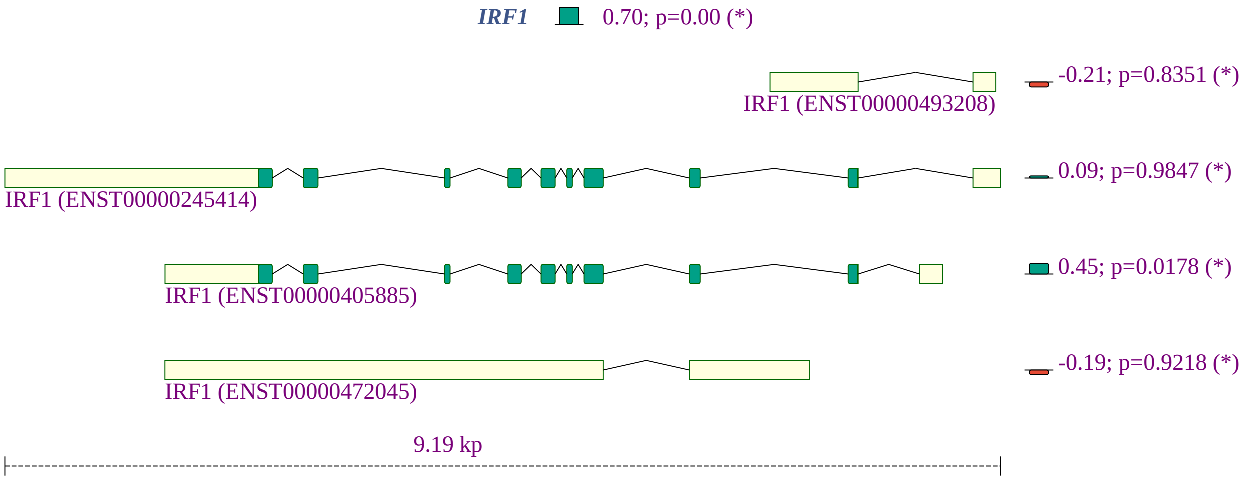


**Figure S5:** *IRF1* isoforms in clinical samples (nasal swabs from individuals with COVID-19). The overall expression of *IRF1* is increased. From top to bottom, ENST00000493208 is a retained intron isoform (nontranslated); ENST00000245414 and ENST00000405885 both encode a protein with 325 residues; ENST00000472045 is a retained intron isoform (nontranslated). The proportion of the protein-coding isoforms increases while that of the retained intron isoforms decreases. The effect of the isoform shift can be predicted to further increase the productive translation of interferon regulatory factor 1.

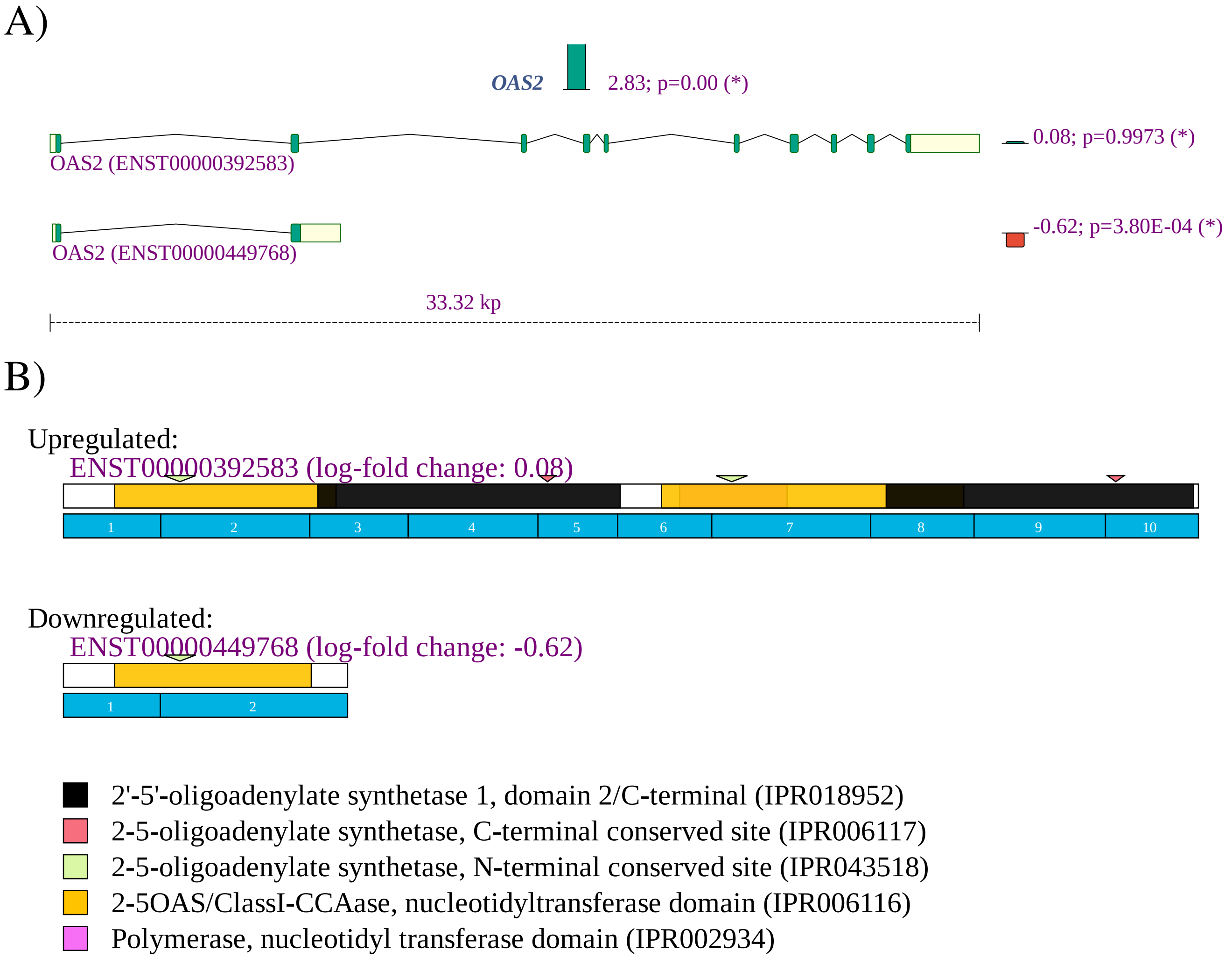


**Figure S6:** *OAS2* **A)** The expression of *OAS2* is increased with a fold change of $2^{2.83} = 7.11$, whereby the expression of a short isoform is reduced. **B)** The short isoform has one OAS domain (UniProt P29728-3), while the long isoform has two (UniProt P29728-2).

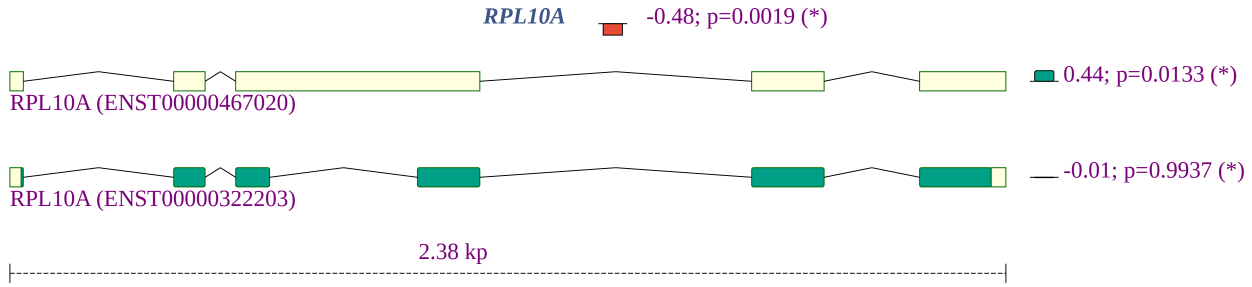


**Figure S7:** *RPLA10A* isoforms in clinical samples (nasal swabs from individuals with COVID-19). The overall expression of *RPLA10A* is decreased and the proportion of the non-coding isoform is increased.

A)

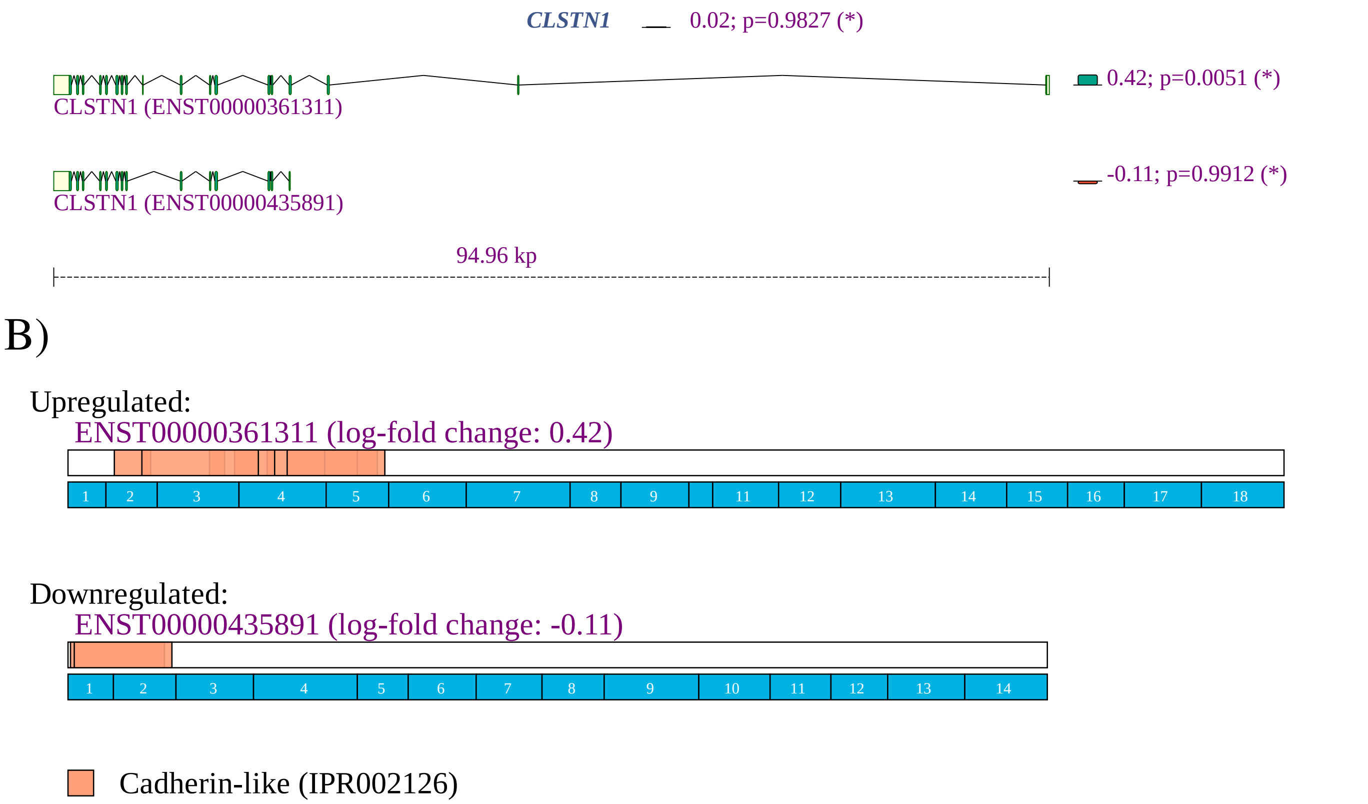


**Figure S8:** *CLSTN1* **A)** The proportion of the longer isoform (ENST00000361311) was increased. **B)** The corresponding protein has two CADHERIN 2 motifs (Prosite PS50268).

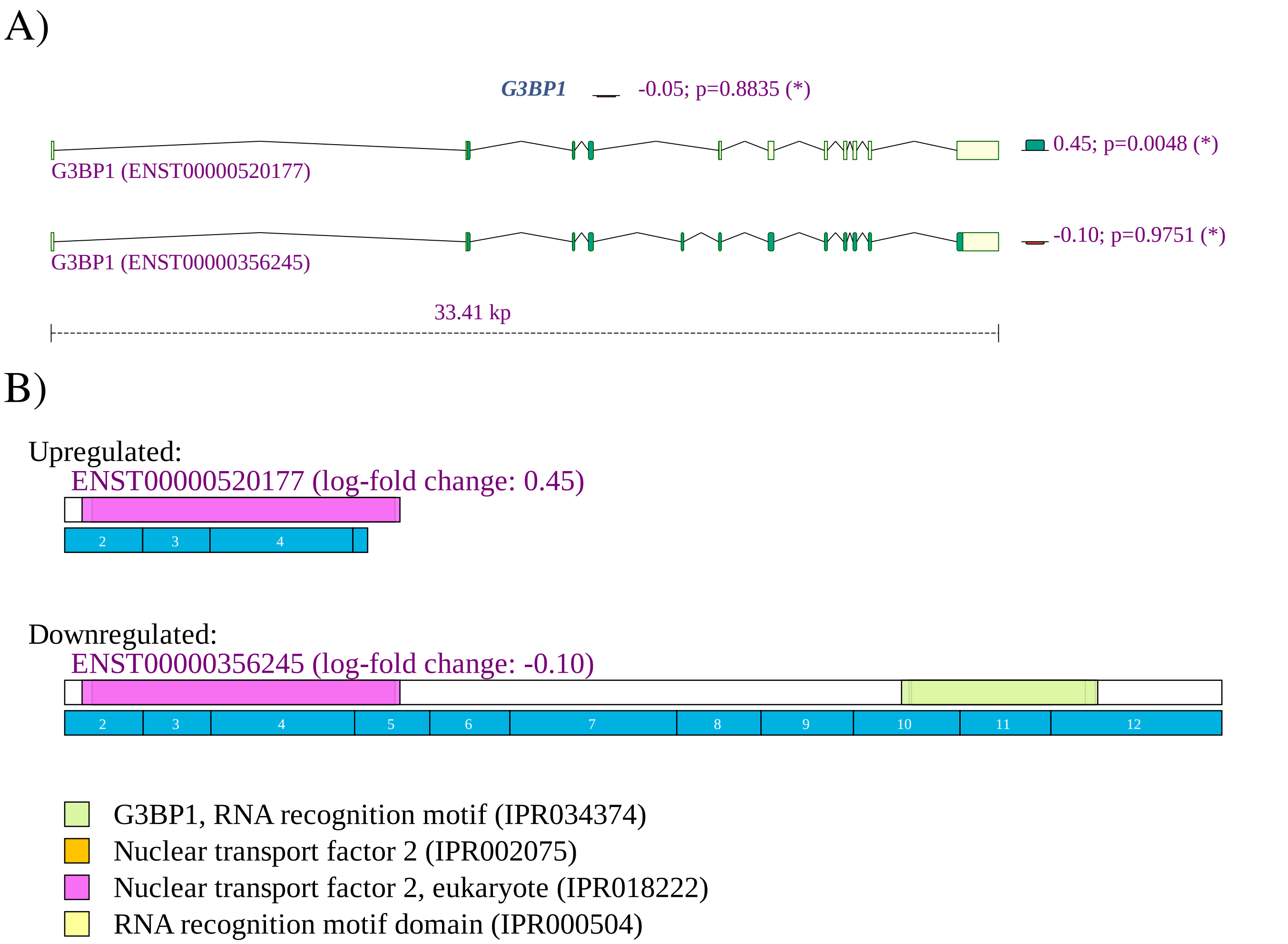


**Figure S9:** *G3BP1* **A)** The proportion of the shorter isoform (ENST00000520177) was increased. **B)** The shorter isoform lacks the RNA recognition motif (RRM) domain (Interpro IPR034374).

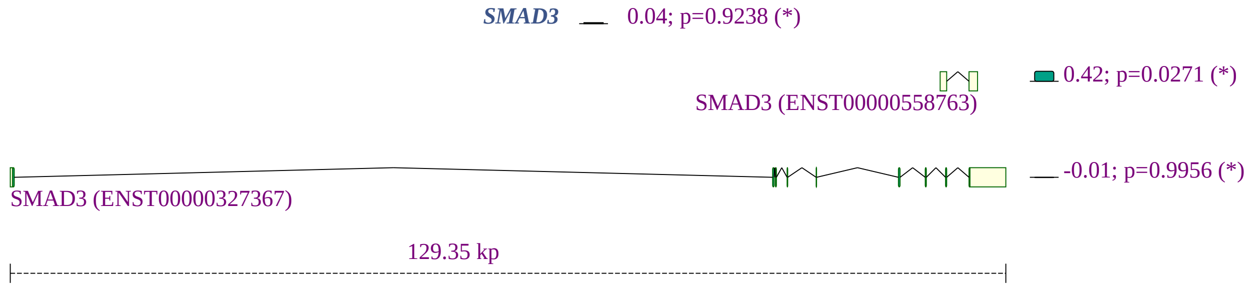


**Figure S10:** *SMAD3* isoforms in clinical samples (nasal swabs from individuals with COVID-19). A shift to the retained intron (non-coding) isoform ENST00000558763.1 is shown.
